# Structural basis for selective inhibition of human GABA transporter GAT3

**DOI:** 10.1101/2025.03.27.645797

**Authors:** Jonas Sigurd Mortensen, Francesco Bavo, Alexander Peder Smiszek Pedersen, Julian Philipp Storm, Tillmann Pape, Bente Frølund, Petrine Wellendorph, Azadeh Shahsavar

**Affiliations:** Department of Drug Design and Pharmacology, Faculty of Health and Medical Sciences, University of Copenhagen, Copenhagen, Denmark; Structural Molecular Biology Group, Protein Structure & Function Program, Novo Nordisk Foundation Center for Protein Research, Faculty of Health and Medical Sciences, University of Copenhagen, Copenhagen, Denmark; Core Facility for Integrated Microscopy (CFIM), Faculty of Health and Medical Sciences, University of Copenhagen, Copenhagen, Denmark

## Abstract

The astrocytic γ-aminobutyric acid (GABA) transporter, GAT3, is essential for terminating GABAergic signalling in the central nervous system. Selective inhibition of GAT3 offers a potential strategy for elevating extracellular GABA levels for the treatment of neurological disorders including epilepsy. However, few potent and selective GAT3 inhibitors have been developed, and their mechanisms of inhibition remain poorly understood. Here, we present the cryo-electron microscopy structures of full-length, wild-type human GAT3 bound to a selective inhibitor, to substrate GABA, or in substrate-free state. GAT3 bound to the inhibitor or in the substrate-free state exhibits an inward-open conformation. The inhibitor binds within the intracellular permeation pathway, positioned between transmembrane helices 1, 2, 3, 6, 7, and 8. The GABA-bound GAT3 is captured in an inward-occluded state, revealing the ion coordination and substrate recognition network, including a cation-π interaction between GABA’s γ-amino group and a phenylalanine residue in transmembrane helix 6. Our data reveal the molecular determinants for the inhibitor selectivity, and the mode of substrate binding and transport inhibition, providing blueprints for the rational design of next-generation selective GAT3 inhibitors.

## Introduction

γ-Aminobutyric acid (GABA) is the main inhibitory neurotransmitter in the central nervous system (CNS). GABA exerts its inhibitory effects via ionotropic GABA_A_ receptors and metabotropic GABA_B_ receptors, leading to hyperpolarization of the postsynaptic membranes. Master keys in the regulation of GABA signalling are the transmembrane GABA transporters (GATs) that belong to the family of secondary active neurotransmitter/sodium symporters (NSSs) of the solute carrier 6 (SLC6) ^1^. The GATs terminate GABA neurotransmission by taking up excess GABA into nearby subcellular compartments. Among the four GAT subtypes (GAT1, GAT2, GAT3 and BGT1; human nomenclature), GAT1 (*slc6a1)* and GAT3 (*slc6a11*) are the most prevalent in the CNS. GAT1, located primarily on neurons, mediates uptake of GABA into presynaptic neurons, thus permitting re-cycling of neurotransmitter, whereas GAT3, expressed solely on astrocytes, governs glial GABA uptake and additionally supports oxidative GABA metabolism^2–4^. The metabolic turnover of GABA into glutamine sustains the downstream bidirectional interaction between neurons and astrocytes and highlights GAT3 as an interesting - yet understudied - target for regulating overall network excitability^5,6^.

Dysregulation of GABAergic signalling with a specific involvement of GAT3 has been implicated in various neuropsychiatric disorders, including epilepsy, Alzheimer’s disease, and ischemic stroke^7–11^. Potentiating GABA-mediated transmission has proved to be an effective approach to suppress seizure activities in epilepsy. Since the approval of tiagabine, a lipophilic, nipecotic acid-derived GAT1 inhibitor, nearly three decades ago, no other GAT inhibitors have been developed for the treatment of epilepsy^12^. Due to its glial localization, pharmacological inhibition of GAT3 is attractive as it offers an alternative therapeutic approach to elevating extracellular and/or extrasynaptic GABA levels, and may thus enhance inhibitory neurotransmission in a manner distinct from GAT1^13–15^. However, very few GAT3 inhibitors have been developed and pharmacologically evaluated, mainly due to a limited potency and selectivity, and, thus far, no GAT3 inhibitors have reached clinical translation.

These limitations, in part, stem from the lack of detailed structural information on the GAT3 binding site, which has led to a simplistic approach in designing GAT3 inhibitors, primarily through N-substitution of known GAT3 substrates with specific bulky lipophilic moieties, conferring GAT3 selectivity over GAT1. Among these inhibitors, (*S*)-SNAP-5114 is the most well-known, but its poor chemical stability restricts the in vivo applications. Inspired by DPPM-1457, a more stable (*S*)-SNAP-5114 variant, we recently developed the GAT3 inhibitor (2*S*, 2’*R*)-2-hydroxy-2-(1-((*E*)-4,4,4-tris(4-methoxyphenyl)but-2-en-1-yl)pyrrolidin-2’-yl)acetic acid, in brief *SR*-THAP^16,17^. Notably, *SR*-THAP is derived from a substrate-like inhibitor that is otherwise nearly inactive.

The process of neurotransmitter uptake by GAT3 and other members of NSS, is coupled to co-transport of Na^+^ (and Cl^−^ in eukaryotic members), where the transporter cycles through distinct conformational states, wherein the substrate-binding pocket is alternately exposed to the extracellular and the intracellular spaces^18–28^. Interestingly, GAT3 may also operate in reverse, contributing to the tonic inhibition of neurons^29^. Detailed structural insights into GAT3 and its interaction with the substrate and inhibitor candidates are essential for understanding the molecular basis of GAT3 uptake and the development of effective pharmacological agents. Here, we present the cryo-electron microscopy (cryo-EM) structures of human GAT3 bound to the selective inhibitor *SR*-THAP, to the substrate GABA, and in the substrate-free state. We capture the inhibitor-bound GAT3 and substrate-free states in an inward-open conformation, while the GABA-bound state is in an inward-occluded conformation. Our study reveals the structural determinants and mechanism of inhibition of GABA transport by GAT3.

## Results

### *SR*-THAP is a selective inhibitor of GAT3

We have designed *SR*-THAP and its enantiomer *RS-*THAP with the aim of developing potent and selective GAT3 inhibitors (Supplementary Fig. 1) ^17^. To evaluate the inhibition mode of *SR*-THAP, we measured GABA uptake inhibition at GAT3 before and after pre-incubation with the inhibitor. Without pre-incubation, *SR*-THAP inhibited the uptake of GABA in mammalian cells (HEK293T cells) expressing human GAT3 with a half-maximal inhibitory concentration (IC_50_) of 25.6 µM, while its enantiomer, *RS*-THAP, showed only limited effect (max 25% inhibition at 300 µM) (Fig. 1a,b). We then measured GABA uptake inhibition at GAT3 after pre-incubation with *SR*-THAP for 120 minutes, which significantly left-shifted the curves, yielding IC_50_ values of 4.9 µM for *SR*-THAP and 140 µM for *RS*-THAP (Fig. 1a,b). The inhibition mode of *SR*-THAP at GAT3 resembles the dual inhibition mechanism of tiagabine at GAT1, which is dependent on pre-incubation conditions, where the inhibitor binds the transporter from both the extracellular and the intracellular sides^23,30^. As expected, the IC_50_ value for the substrate GABA remained unchanged also after preincubation, as the substrate binds the transporter solely from the extracellular space (Fig. 1c).

**Figure 1.**
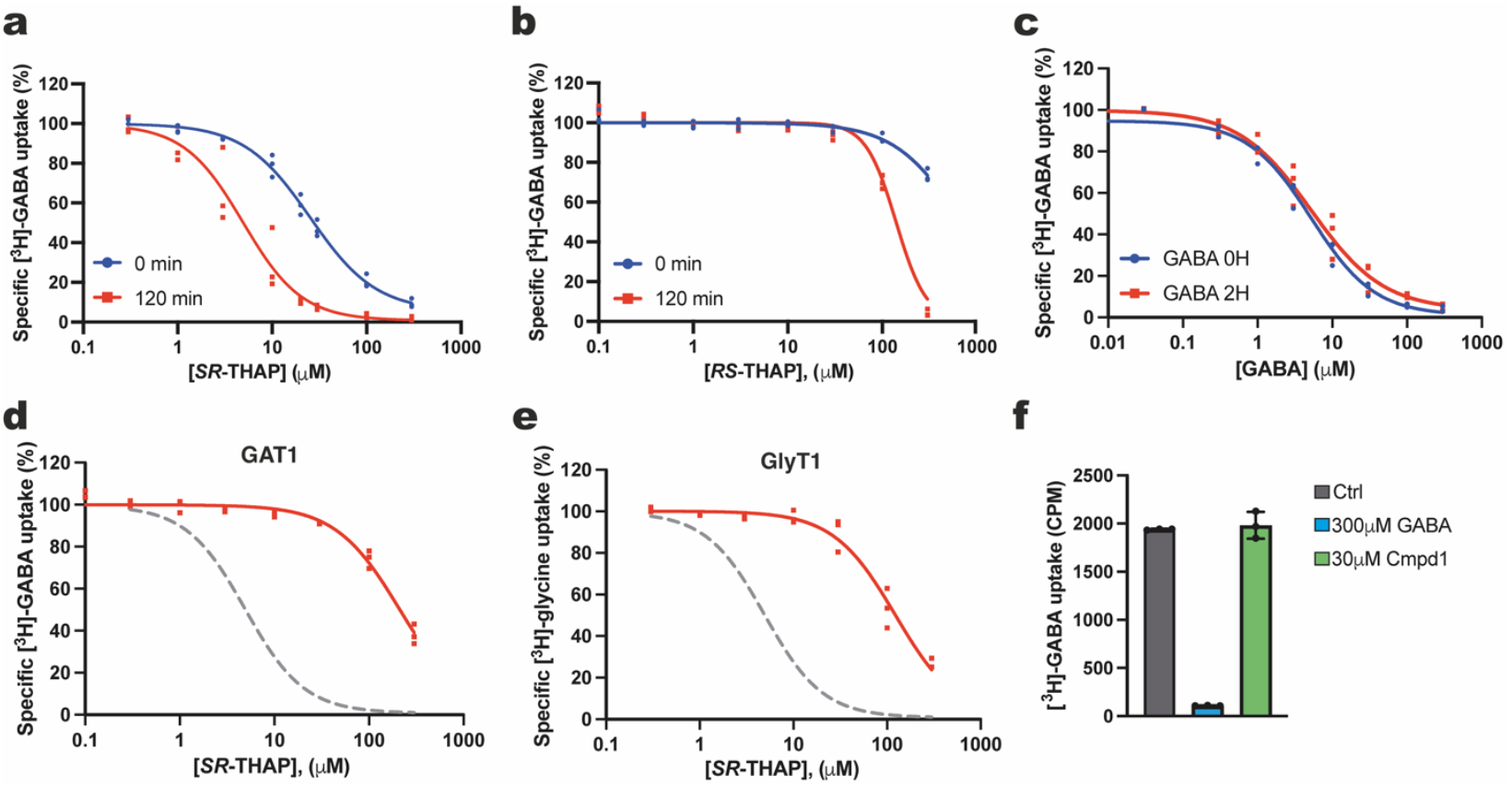
*SR*-THAP is a selective inhibitor of GAT3 with a dual inhibition mechanism. **a**, *SR*-THAP inhibits [^3^H]GABA uptake by GAT3 expressed in HEK293T cells with an IC_50_ of 25.2 [21.8; 30.4] µM without pre-incubation (blue) and 4.9 [3.2; 7.0] µM with 120 min pre-incubation. **b**. The enantiomer *RS*-THAP also inhibits [^3^H]GABA uptake by GAT3 expressed in HEK293T cells, but with significantly reduced potency, estimated IC_50_ values of >600 µM without pre-incubation and >100 µM with pre-incubation. **c**, [^3^H]GABA uptake by GAT3 can also be inhibited by non-radioactive GABA, although with minimal observed shift in potency comparing with and without pre-incubation of 120 min, with IC_50_ values of 8.5 [6.4; 11] and 5.4 [4.6; 6.4] µM respectively. **d**, The GlyT1 specific benzoylisoindoline inhibitor Cmpd1 has no effect on GAT3 mediated [^3^H]GABA transport even at 30 µM. **e**, Inhibition of [^3^H]GABA uptake by GAT1 in HEK293T cells by *SR*-THAP did not reach full inhibition at the highest concentration tested (300 µM); estimated IC_50_ of 212 [143; 238] µM. **f**, Similarly to GAT1, inhibition of [^3^H]glycine uptake by GlyT1 in HEK293T cells did not reach full inhibition at 300 µM; estimated IC_50_ of 123 [104; 147] µM. Data points are shown as individual replicates from n = 3 biological replicates (triplicate measurements) unless othervise stated.

We then investigated the selectivity of *SR*-THAP, given its higher potency compared to its enantiomer. *SR*-THAP was found to be respectively 42 and 23 times more selective for GAT3 over GAT1 (estimated IC_50_ value of 212 µM) and GlyT1 (estimated IC_50_ value of 117 µM) (Figure 1 d-f). Further, the high-affinity benzoylisoindoline inhibitor of GlyT1, Cmpd1^24,31^, exhibited no effect on GAT3-mediated GABA uptake at concentrations up to 30 µM (approximately 2300 times the IC_50_ of hGlyT1).

Given the lack of detailed experimental information on the interaction network of *SR*-THAP with GAT3, and to understand the mechanism underlying the inhibition of GABA transport, we next sought to determine the structures of GAT3 bound to the inhibitor and the substrate GABA.

### Overall architecture of human GAT3

We purified wild-type human GAT3 in a glyco-diosgenin (GDN) micelle, obtaining a monodisperse and homogenous sample for cryo-EM studies (Supplementary Fig. 2). We measured GABA uptake nhibition by wild-type un-tagged GAT3 and the cryo-EM GAT3 construct (with affinity tags) and obsereved no difference in IC_50_ values (6.9 µM and 7.0 µM, respectively) (Supplementary Fig. 2). We determined the structures of GAT3 bound to inhibitor *SR*-THAP, substrate GABA, or in substrate-free state at 2.9, 3.2, and 3.9 Å resolution, respectively (Figure 2, Supplementary Figs. 3-7).

**Figure 2.**
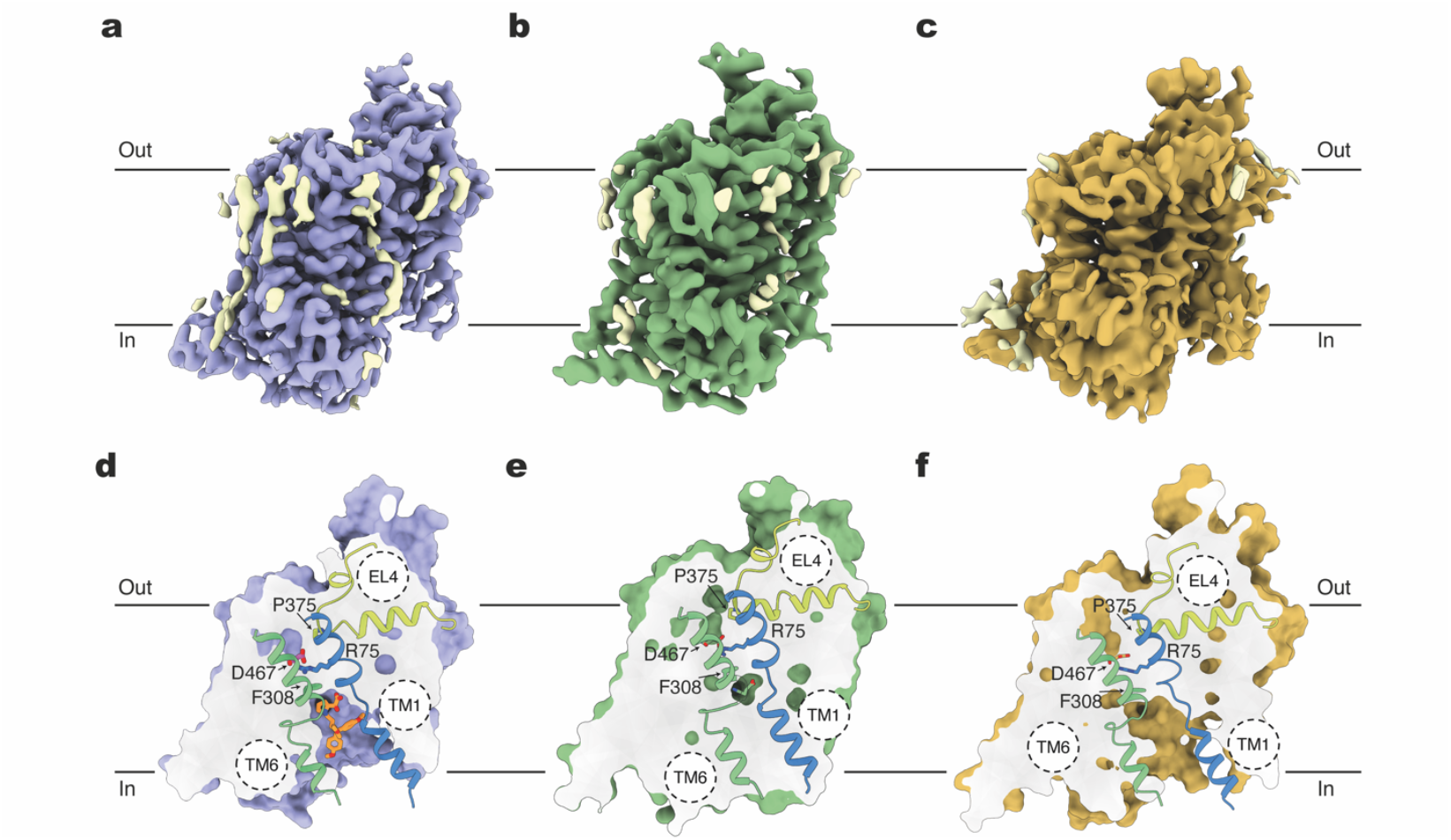
Inhibitor-bound, GABA-bound and substrate-free states of GAT3. Cryo-EM density maps of GAT3 bound to **a**) *SR*-THAP (blue map, contour level = 0.063 in ChimeraX), **b**) GABA (green map, contour level = 0.06 in ChimeraX) and **c**) in the substrate-free form (light orange map, contour level = 0.03 in ChimeraX). Lipid densities are shown in light yellow. **d, e, f**, Surface representation of the inhibitor-bound, GABA-bound and substrate-free structures of GAT3 viewed parallel to the membrane. **d**, Slice view of GAT3 showing the *SR*-THAP (orange) binding pocket. **e**, Slice view of GAT3 showing the GABA (green) binding pocket. **f**, Slice view of GAT3 in the substrate-free state. The extracellular gating residues R75 (TM1 – blue) and D467 (TM10 – not shown), as well as P375 (EL4 – yellow) and F308 (TM6 – green), are shown as sticks in **d, e, f**.

We unambiguously modelled all transmembrane (TM) helices with clear side chain densities, intracellular and extracellular loops except parts of extracellular loop 2 (EL2) due to its high degree of flexibility (Fig. 2, Supplementary Figs, 5-7). The conserved NSS disulfide bridge between C171 and C180 (EL2) is observed in both the inhibitor-bound and substrate-free structures of GAT3. However, in the GABA-bound state, while the densities for C171 and C180 are clearly resolved, we do not observe distinct density for the disulfide bridge, suggesting that the cysteine residues may be reduced in this sample (Supplementary Figs. 5-7, 8, 9).

GAT3 adopts the expected architecture of SLC6 transporters, comprising 12 α-helical TMs with an inverted pseudo-twofold symmetric organisation of TMs 1–5 relative to TMs 6–10, denoted as the LeuT fold^32^, with TMs 1 and 6 unwinding approximately halfway across the membrane, a characteristic for the central binding site in NSS (Fig. 2 and Supplementary Figs 5-7).

Unique to GABA transporters, as well as to taurine and creatine transporters, is an insertion of an extra serine residue, (S472 in GAT3) in the unwound region of TM10 that forms a one-turn π-helix and provides extra bulk in this region. This extra residue has been shown to be required for the coupling of ions for the transport of the substrate GABA^33,34^. In all three GAT3 structures, the side chain hydroxyl group of S472 forms hydrogen bonds with the hydroxyl group of T304 (TM6a) and Arg75 (TM1b), as part of the interaction network occluding the extracellular pathway (Fig. 2, Supplementary Fig. 9). In higher resolution maps of inhibitor-bound and GABA-bound GAT3, we obsereve a clear density for a water molecule that mediates an interation between S472 (TM10) and D301 (TM6a) similar to those observed for inward-facing structures of GAT1 bound to inhibitors and substrate^23,35, 28^.

### *SR*-THAP locks GAT3 in an inward-open conformation

The inhibitor *SR*-THAP captures GAT3 in an inward-open conformational state, where the central substrate site is accessible to the solvent via the intracellular vestibule. The N-terminal segment of TM1 (TM1a) is bent away from the core of GAT3, opening the cytoplasmic release pathway and allowing the accommodation of the bulky inhibitor (Supplementary Fig. 10). The splayed motion of TM1 disrupts the interaction between the TM1a and the hydrophobic patch at the cytoplasmic part of GAT3 that is otherwise present in other outward-facing or inward-occluded structures of NSS^19,21,36–38^. We could not model the intracellular part of TM5 preceding the conserved helix-breaking G(X9)P motif (G251(X9)P261 in GAT3), which we assume to be unwound, a feature that creates a solvent pathway at the intracellular side of the transporter, providing water access from the cytoplasm to the Na2 site^37^.

The highly ordered interaction network that closes the extracellular pathway includes residues from TMs 1, 3, 6, and 10 as well as EL4 (Fig. 2). The conserved extracellular gate residues R75 (TM1b) and D467 (TM10), form a salt bridge with a Cα-Cα distance of 9.6 Å, precluding solvent access from the extracellular side. In addition to this salt bridge, R75 forms a stacking and cation-π interaction with the phenyl ring of the conserved F308 (TM6a) residue. F308 which sterically blocks the extracellular pathway, is within Cα-Cα distance of 11.6 Å to the conserved Y147 (TM3), consistent with previous inward-facing NSS structures^23,24,38–40^.

R75 which is conserved across NSS is critical for the transport of the substrate. The conservative mutation of the equivalent residue in hGAT1, R69K, impairs GABA transport, while sustaining the ability of the tranporter to bind the substrate^41^, indicative of a locked outward-facing conformation of the transporter. The R75-coordinating residues, Q305 (TM6a), D467 (TM10), and S472 (unwound region of TM10), form an extensive hydrogen bond network. Maintaining the integrity of this network is critical for transporter function, as demonstrated by conservative mutations in hGAT1^42,43^.

We observe a clear density for a Cl^−^ ion that is coordinated by the conserved residues Y192 (TM2), Q305 (TM6a), S309 (TM6a) and S345 (TM7) (with mean coordination distance of 3.2 Å ± 0.4) (Supplementary Fig. 9). We do not, however, observe any densities for sodium ions at Na1 or Na2 sites, consistent with an inward-open conformational state of the transporter.

### *SR*-THAP binding mode

The high-quality of the cryo-EM density for *SR*-THAP allowed us to unambiguously model the inhibitor sandwiched in between TMs 1, 2, 3, 6, 7 and 8 (Fig. 3). Each of the three moieties of *SR*-THAP (R^1^-R^3^, Fig. 3) is accommodated in a defined sub-pocket of GAT3, resulting in a modular binding mode similar to that of tiagabine at GAT1^23,35^.

**Figure 3.**
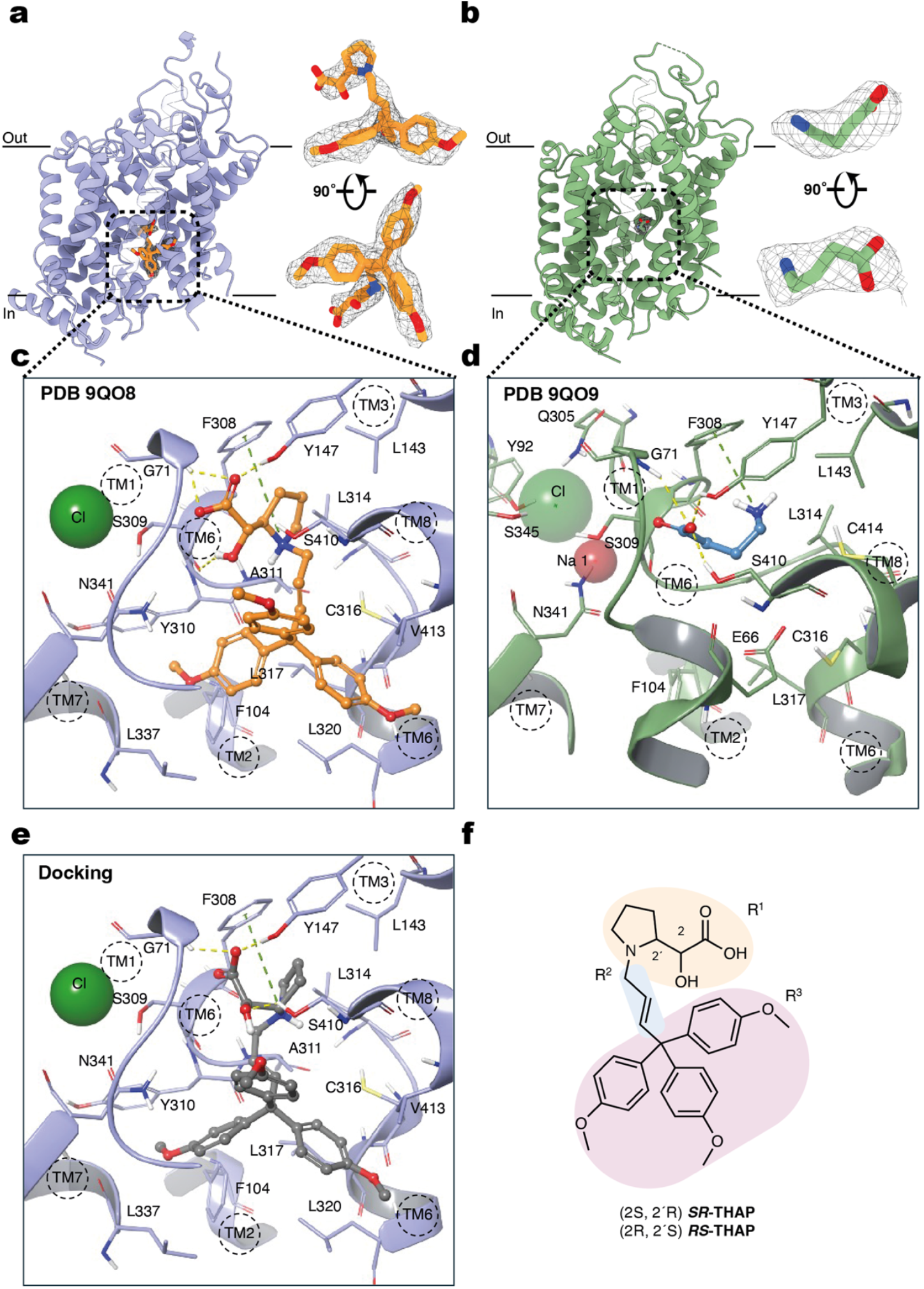
Inhibitor *SR*-THAP and substrate GABA bind to the central binding pocket of GAT3. **a**,**b** Overall structure of the human GAT3 bound to the selective inhibitor *SR*-THAP (orange stick) (**a**), and its substrate GABA (green stick) (**b**). Cryo-EM density of *SR*-THAP (map contour level = 0.072 in ChimeraX) (**a**) and GABA (map contour level = 0.06 in ChimeraX). **c, d, e**, Close-up view of the binding mode of *SR*-THAP (orange) to GAT3 (**c**), the binding mode of GABA (blue-grey) to GAT3 (**d**), and the docking pose of *RS*-THAP (dark grey) to GAT3 (**e**). Protein backbone is shown as ribbons cartoon (azure), chloride ion as green sphere, sodium ion as red sphere and interacting residues within 3 Å are displayed in grey. H-bonds and cation-” interactions are displayed with yellow and green dashed lines, respectively. **f**, Chemical structure of *SR*-THAP and of its enantiomer *RS*-THAP. The three structural elements forming the inhibitor are highlighted in faded orange for the amino acidic moiety R^1^ (2-hydroxy-2-pyrrolidin-2-yl)acetic acid); in faded azure for the N-alkyl chain R^2^; in faded pink for the trimethoxytrityl group R^3^.

The inhibitor binding site is capped by the aromatic side chains of F308 (TM6a) and Y147 (TM3) as well as G71 (TM1), that together with the lateral L143 (TM3) form a lipophilic cavity where R^1^ (the amino acidic moiety) is lodged. This cavity is further lined by A301 and L314 from the non-helical region of TM6. The residue L314 is part of the conserved (G/A/C)ΦG motif (C_313_L_314_G_315_ in GAT3), which plays a central role in shaping the binding site to accommodate ligands of varying sizes across the SLC6 family^44^. A conserved glutamate residue in TM2, E107 in GAT3, is coordinating the CLG loop through a direct and a water-mediated hydrogen bond with backbone nitrogen of G315 (Supplementary Fig. 9). This ordered water molecule is also observed in previous NSS structures including GAT1 structure bound to substrate GABA (PDB ID 7Y7W) ^35^. The interaction between E107 (TM2) and the non-helical part of TM6 is conserved among SLC6 and appears as a key functional element of transport^45^. Mutagenesis of the equivalent glutamate residue in GAT1 diminishes the transport activity, supporting its important role in a functional transporter^46^.

The positively charged nitrogen of R^1^ establishes a cation-*π* interaction with F308, while the hydroxyl group of R^1^ interacts via a hydrogen bond with the amidic carbonyl of S309 (TM6). The carboxylate group of R^1^ extends toward TM1 and interacts with hydroxyl of Y147 (TM3) as well as the amidic NH of L70 and G71 (TM1), similar to that of substrate-transporter interactions, in *α*-amino acid transporter structures^24,37,44,47^. The carboxylate of R^1^ further interacts with TM8, through water mediated hydrogen bonds with L406 and S410.

The R^2^ moiety (N-alkyl chain) elongates into a narrow lipophilic opening toward the bottom of the binding site, and it is surrounded by the side chains of L314 (CLC motif on TM6), A311 (TM6), S410 (TM8), and V413 (TM8). The R^3^ group (trimethoxytrityl moiety) is accommodated within a lipophilic cavity at the bottom of the binding site, exposed to the cytosol, shaped as a tripod, and defined by residues from TMs 1, 6, 7 and 8 – G66, P104, Y310, C316 (CLC motif in TM6), L320, L337, L406, V413 and N431.

Molecular docking places *RS*-THAP, DPPM-1457, and (S)-SNAP-5114 in the binding pocket of GAT3, with (S)-DPPM-1457 and (S)-SNAP-5114 matching the binding mode of *SR*-THAP, with preserved hydrogen bonds between the carboxylate groups and G71 and Y147 residues (Fig. 3, Supplementary Fig. 11). The R^2^ and R^3^ groups of the less active enantiomer of *SR*- THAP – *RS*-THAP – superpose well with that of *SR*-THAP. However, the inversion of the stereocenters in R^1^ distorts its positioning and disrupts the hydrogen bond network, causing the carboxylate and hydroxyl groups of *RS*-THAP to orient differently from those in *SR*-THAP, and the R^2^ group to twists toward TM6.

Comparing the binding-pocket residues of *SR*-THAP -bound GAT3 with corresponding residues in tiagabine-bound GAT1 (PDB IDs 7Y7Z and 7SK2) points to structural determinant for selectivity of *SR*-THAP for GAT3 against GAT1. The small polar E66 in GAT3 (TM1) is replaced by a large aromatic residue, Y60 in GAT1, reducing the size of the vestibular lipophilic sub-pocket and precluding access to R^3^ of *SR*-THAP (Supplementary Fig. 10).

### GABA-bound GAT3 is in an inward-occluded state

GABA is bound in the central cavity of GAT3 formed by residues from TMs 1, 3, 6, and 8, midway through the membrane bilayer. The GABA binding pocket is inaccessible from both the extracellular and intracellular sides, indicative of an occluded state of the transporter. The extracellular gate is closed by the conserved salt bridge between R75 (TM1b) – D467 (TM10) (C-C distance of 9.3 Å), and the intracellular pathway is closed by TM1a (Fig. 2). The distorted density observed in the intracellular half of TM5 (residues 246 – 253) suggests an unwound region at the conserved G251(X9)P261 motif, leading to the solvation of the Na2 site, and indicative of an inward-occluded state of GABA-bound GAT3.

The carboxylate group of GABA forms hydrogen bonds to backbone nitrogen atoms of G71 (TM1) L70 (TM1), and to the hydroxyl groups of the conserved Y147 on TM3 and S410 on TM8. The carboxylate also coordinates the Na^+^ ion with a prominent 10.0 r.m.s.d. signal at the Na1 site. The coordination of Na1 in GABA-bound GAT3 is similar to that in previous NSS structures showing octahedral geometry by side chains of N72 (TM1), N341 (TM7) and S309 (TM6), backbone carbonyl of I67 (TM1) and S309 (TM6), in addition to the GABA carboxyl group. The binding mode of the carboxylate group of GABA resembles that of substrate- and inhibitor-bound structures of NSS with amino acid-based components such as *SR*-THAP-bound GAT3 and GABA-bound GAT1, and in *α*-amino acid transporters such as GlyT1, and bacterial homologues of NSS, LeuT and MhsT^28,35–37,47–49^.

While the carboxylate group interactions of GABA in GAT3 are similar to those in GABA-bound GAT1, the positioning of the amino group differs between the two transporters. In GAT3, the amino group of GABA engages in a cation-π interaction with the phenyl ring of F308 (TM6), whereas in GAT1, it adopts a different orientation, forming a hydrogen bond with the hydroxyl group of Y60 (TM1) ^35^.

Residues involved in substrate recognition among GABA transporters are highly conserved (Supplementary Fig. 3). However, Y60, which interacts with GABA in GAT1, is substituted by a glutamate residue in GAT2, GAT3 (E66), and BGT1. Mutation of this tryrosine residue to glutamate in GAT1 has been shown to abolish the transport activity^41^. In our GABA-bound GAT3, E66 is too far from the substrate to contribute to GABA recognition. Further, the quality of density in the region containing E66 is high, with side-chain densities clearly resolved for all neighboring residues but E66, suggesting significant mobility of this residue. The binding pocket of GABA is further lined by the conserved (G/A/C)ΦG motif (C_313_L_314_G_315_ in GAT3) on the unwound region of TM6^44^.

In addition to the Na^+^ density in Na1 site, we observe a clear density for the Cl^−^ ion (a prominent 14.0 r.m.s.d. signal) that is coordinated by the same set of residues as in the inhibitor-bound GAT3, Y192 (TM2), Q305 (TM6a), S309 (TM6a) and S345 (TM7) (with mean coordination distance of 2.8 Å ± 0.1) (Supplementary Fig. 9). We do not observe any sodium ion density at the Na2 site, consistent with its solvation caused by unwinding of intracellular half of TM5.

### Inward-open state of substrate-free GAT3

Using 3D classification on the *SR*-THAP-bound GAT3 data set, we could identify a subclass where we were not able to observe any density corresponding to the inhibitor. We present this structure as a substrate-free state of GAT3, which is captured in an inward-facing open conformational state. Similar to other inward-open NSS structures, TM1a has splayed away from the transporter core, opening the intracellular permeation pathway. In substrate-free GAT3, the angular movement of TM1a closely resembles that in *SR*-THAP-bound GAT3. However, TM1a in substrate-free state is 6 degrees more open compared to *SR*-THAP-bound form. This was measured using the Cα atoms of G69 on TM1 as the fixed reference point and V57 on TM1a of the two structures as the moving points. In the substrate-free state of GAT1 (PDB ID 7Y7V) ^35^, TM1a is positioned further away from the transporter core, with a 12-degree difference compared to the substrate-free GAT3. The intracellular part of the TM5 is unwound, similar to inhibitor-bound state of GAT3. We do not observe any density corresponding to the Cl^−^ or Na^+^ ions, as expected for an inward-open transporter. However, given the limited resolution of this state, the absence of the ions cannot be definitively confirmed.

## Discussion

We have resolved the first structures of wild-type human GABA transporter GAT3 bound to the selective inhibitor *SR*-THAP, to substrate GABA, as well as in a substrate-free state, without using fiducial markers. We reveal that the inhibitor binds at the intracellular permeation pathway, sandwiched in between TMs 1, 2, 3, 6, 7 and 8, and locks the transporter in an inward-open state (Fig. 4). This cavity remains open in the inward-open substrate-free state of GAT3 where TM1a is by 6 degrees more open compared to the *SR*-THAP-bound structure. The GABA-bound state of GAT3 is captured in an inward-occluded state. The interaction netwrok stabilising GABA in the binding site, closely resembles those observed in eukaryotic and prokaryotic *α*-amino acid transporters of SLC6 family as well as in GAT1, with one main difference. In GAT3, The γ-amino group of GABA interacts with TM6 through a cation-π interaction with the phenyl ring of F308, whereas it interacts with TM1 through a hydrogen bond (Y60) in GAT1. Even though the central substrate binding pocket is similarly surrounded by TMs 1, 3, 6 and 8 across NSS members, the binding modes of biogenic amine serotonin and dopamine transporters are different from that of GAT3 and GAT1^50,51^. In all three cytosol-facing GAT3 structures, the intracellular part of the TM5 is unwound allowing solvation of the Na2 site and the intracellular release pathway.

**Figure 4.**
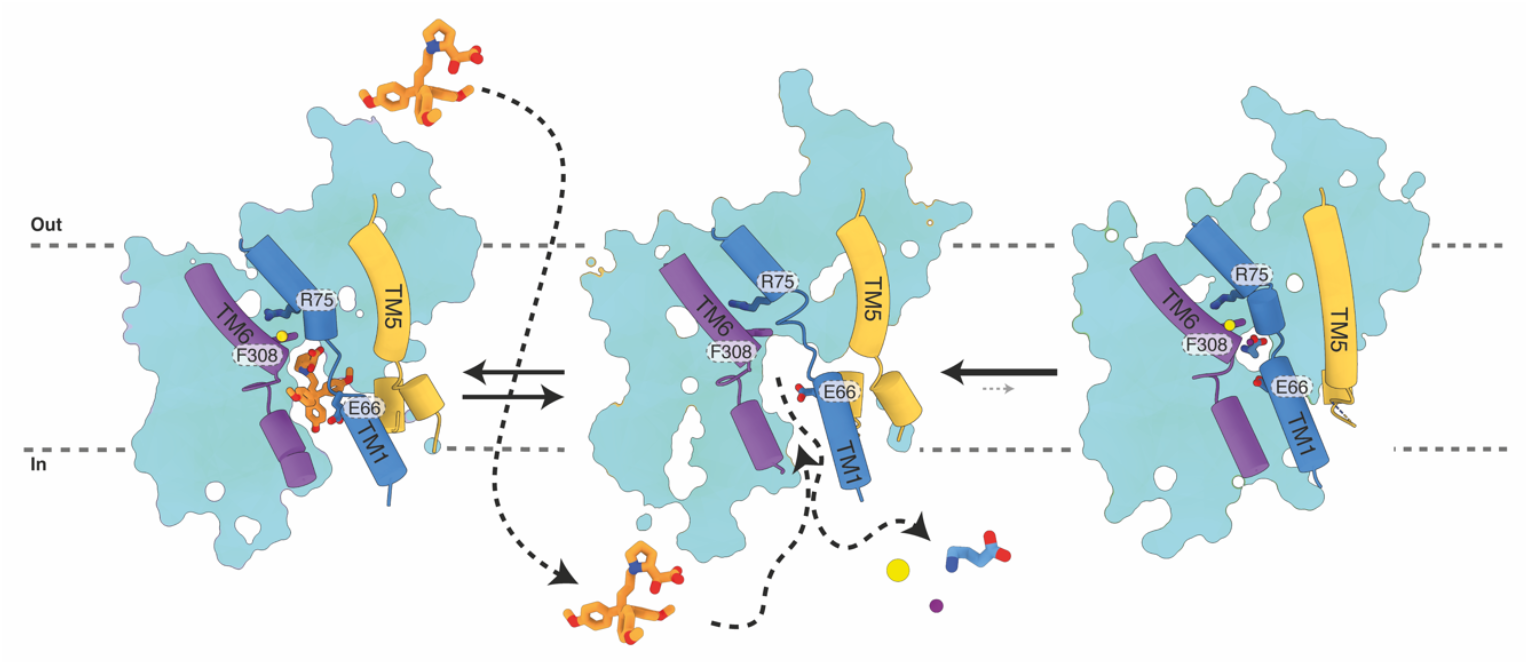
Schematic illustration of the inward facing states of GAT3 the transport cycle. Occlusion of the extracellular pathway is maintained by a highly conserved interaction network including a salt bridge between R75 and D467 (not shown) and stacking and cation-π interaction of R75 and F308. From the occluded GABA-bound state, the transporter transitions to an inward-open state, by an outward tilt of TM1a as well as partial unwinding of the intracellular part of TM5. The GAT3 selective inhibitor *SR*-THAP, after diffusion across the plasma membrane, binds GAT3, locking the transporter in an inward-open conformation. The residue E66 in the binding pocket of GAT3, is replaced by a large aromatic tyrosine in GAT1, playing a part in defining the selectivity of *SR*-THAP for GAT3, against GAT1. Cl^−^ and Na^+^ ions are shown as yellow and purple spheres.

Traditionally, within drug discovery of GABA transporters, the focus has been on GAT1, highly encouraged by the clinical usefulness of the inhibitor tiagabine^12^. However, for enhancing GABAergic signalling, GAT3 inhibition offers a therapeutic advantage over GAT1. GAT3 inhibition not only increases synaptic GABA levels but concomitantly leads to a decrease in astrocytic GABA oxidative metabolism^52,53^. Proof-of-principle for targeting GABA metabolism, e.g. in epilepsy, is evident from the clinical effects of vigabatrin, a blocker of GABA metabolism which increases GABA brain levels^54^. Although experimental GAT3 ligands have been found to display anticonvulsant properties in some preclinical models of epilepsy^7,55^, only few studies are reported, largely due to the limited availability of selective and potent tool compounds. In addition, the most widely used ligand, (*S*)-SNAP-5114, while modestly potent and selective for GAT3, has significant liabilities that not only complicate the interpretation of its efficacy per se, but also limit insights into GAT3 as a therapeutically relevant drug target. These liabilities include poor chemical stability^16^, limited solubility, limited brain uptake after systemic administration^55^, reports of cellular toxicity^9^ as well as severe in vivo toxicity when tested in mice with stroke^56^. Thus, the lack of GAT3-selective tool compounds has hampered the evaluation of GAT3 as a relevant drug target, and the absence of structural insight has vice versa limited the design of better inhibitors.

In this study we present the binding mode of a novel inhibitor *SR*-THAP which is superior to (*S*)-SNAP-5114 in terms of chemical stablity and potency and with improved selectivity at GAT3 over GAT1 compared to DPPM-1457. *SR*-THAP is the result of a medicinal chemistry design proces which can now be rationalized further based on the cryo-EM data presented here. *SR*-THAP was designed together with *RS*-, *SS*- and *RR*-THAP as an N-substituted analogues of small and conformationally constrained derivatives of (*S*)-isoserine. As *N*- substituent, we chose the (*E*)-1-4,4,4-tris(4-methoxyphenyl)but-2-en-1-yl group of DDPM-1457. To our surprise, compound *SS*-THAP was inactive even though it was derived from the only active analogue of (*S*)-isoserine. Conversely, *SR*-THAP and its enantiomer *RS*-THAP displayed inhibition of hGAT3-mediated GABA uptake, even though they were derived from the inactive precursors^17^. Thus, despite starting from a nearly inactive substrate, in which N-substitution alone would not typically be expected to introduce activity^57^, the strategic incorporation of the bulky and stable lipophilic moiety from DPPM-1457 surprisingly reintroduced significant GAT3 potency (IC_50_ (2h) of 4.9 µM) ^17^. This suggests that this type of modification may not only enhance potency but also maintain or improve selectivity.

To rationalize these findings, we hypothesized that small amino acid-based and bulky *N*-substituted hGAT3 inhibitors could exhibit different binding modes at GAT3. Inspired by the recent cryo-EM structures of tiagabine-bound GAT1^23,35^, we further hypothesized that *SR*-THAP and structurally related inhibitors could bind GAT3 at a similar intracellular binding site. Therefore, we took a two-thonged approach aimed at i) investigating the compound mode of inihibition of *SR*-THAP, and ii) providing structural evidence of the binding mode.

In pharmacological uptake studies at GAT3, a 2-hr preincubation with *SR*-THAP demonstrated a 10 times shift in potency, suggesting that the compound may need to permeabilize the cell membrane to access its binding site. Based on the high-resolution structure *of SR*-THAP-bound GAT3, we now provide the structural evidence for a less conserved binding site at the inward open facing conformation of GAT3.

These findings will allow structure-based rational design of novel potent and selective GAT3 inhibitors and next-generation therapeutics. Since this novel binding site is less conserved than the central pocket of SLC6 transporters, it offers an opportunity for the generation of more selective compounds. Additionally, the strict stereochemical requirements highlighed by the inactive enantiomer *RS*-THAP may further inform drug design. While the ligand-receptor shape complementarity within the non-conserved, tripod-shaped sub pocket (R^3^) introduces GAT3 activity in otherwise weakly active amino acids^57^, an appropriate hydrogen bond network between the amino acid core and the more conserved sub pocket (R^1^) is crucial^17^. Moreover, maintaining a relatively rigid and narrow spacer (R^2^) ^58^ is essential for preserving GAT3 potency. Future GAT3 inhibitors may not only offer therapeutic potential for epilepsy, but could also be highly relevant tool compounds for studying conditions that involve aberrant GABA neurotransmission, such as stroke^9^, Alzheimer’s disease^11^, and neuroinflammation^8^.

## Data availability

Atomic coordinates of GAT3 bound to *SR*-THAP, GABA, or in substrate-free state have been deposited in the Protein Data Bank (PDB) under accession codes 9QO8, 9QO9, and 9QOA, respectively. The corresponding cryo-EM maps have been deposited in the Electron Microscopy Data Bank (EMDB) under accession numbers EMD-53259, EMD-53260, and EMD-53261, respectively.

## Acknowledgements

We thank Ryan P. Cantwell Chater for helpful discussions on cryo-EM data processing. The plasmid for expression of His-tagged HRV-3C protease was a kind gift from Dr. Eric R. Geertsma. This project has received funding from the Lundbeck Foundation (R368-2021-522) to A.S., and the Novo Nordisk Foundation (NNF21OC0067835) to P.W, and NFF20OC0065017 to B.F., and the Independent Research Fund Denmark (1026-00335B) to P.W. We thank L. Kristensen for access to the protein production facility at the Department of Drug Design and Pharmacology, University of Copenhagen. The cryo-EM data was collected at the at the Danish Cryo-EM Facility at the Core Facility for Integrated Microscopy (CFIM), University of Copenhagen, supported by the Novo Nordisk Foundation (NNF17SA0024386, NNF22OC0075808, and NNF24SA0098829). We acknowledge the expert support offered by Nicholas Sofos at CFIM.

## Contributions

J.S.M., P.W., and A.S. conceived and designed the project. F.B. and B.F. developed and provided *SR*-THAP compound. J.S.M. and A.S. established the protein expression and purification conditions. Expression, purification, and cryo-EM sample preparations were carried out by J.S.M. with support from A.S. Data collections, processing, analysis and structure determination were carried out by J.S.M. with assistance from J.P.S., T.P., and A.S. GABA uptake experiments were carried out by A.P.S.P. with assistance from J.S.M. Molecular docking was carried out by F.B. J.S.M., F.B., P.W., and A.S. wrote the initial draft of the manuscript with contributions from all authors.

## Methods

### GAT3 constructs and cell lines

The human GAT3 complementary DNA sequence was codon optimised and synthetised by GenScript for expression in mammalian cells. For protein expression in Expi293F cells, a construct carrying the sequence of GAT3 followed by a G(GGGS)_3_G linker, a HRV 3C protease cleavage site (LEVLFQGP), enhanced green fluorescent protein (eGFP), a twin Strep-Tag II tag and a decahistidine tag were cloned into a pCDNA3.1 vector. This construct is referred to as GAT3_Cryo_ in supplementary figure 2. Furthermore, for uptake experiments in HEK293T cells, wild-type human GAT3 and GAT1 constructs previously generated (hGAT3 FRT pcDNA5 and hGAT1 FRT pcDNA5) were used^59^ or wild-type human GlyT1 carrying an eleven amino acid HiBit tag at the c-terminal end (hGlyT3-HiBit pcDNA3). All constructs were verified by DNA sequencing. The Expi293F cell stock was purchased from ThermoFisher Scientific and confirmed to be mycoplasma-free. The cell line was regularly tested for mycoplasma contamination and tested negative for contamination. No misidentified cell lines were used. The HEK-293T cell line was purchased from ATCC (293T-ATCC; #CRL-3216; authenticated to be mycoplasma-free).

### Protein expression and purification

GAT3 was expressed in HE400AZ expression medium (GMEP Cell Technologies, Japan) in 600 mL TubeSpin bioreactors, incubating in an orbital shaker at 37°C, 8% CO_2_ and 175 rpm in a humidified atmosphere. The Expi293F cells were transfected at a density of 3.0 – 3.5 × 10^6^ cells per mL and a viability of above 97%. A 25 kDa linear polyethylenimine (LPEI) was used as the transfection reagent, at a GAT3 DNA:LPEI ratio of 1:1 with a total plasmid concentration of 2.0 µg per mL of culture volume to transfect. The LPEI/DNA complexes were incubated in Opti-MEM™ I Reduced Serum Medium (ThermoFisher Scientific) for 15 minutes before adding to the cells. At approximately 20 hours post-transfection, cell cultures were supplemented with 10 mM sodium butyrate. The cells were harvested at approximately 68 hours post-transfection at a viability of above 70% and stored at −70 °C until purification.

The cell pellets were thawed on ice and resuspended in 50 mM Tris-HCl (pH 7.5), 150 mM NaCl, 1 cOmplete™ Protease Inhibitor Cocktail tablet per 50 mL of buffer (Roche), 0.25 mg of DNAse I and incubated on rotation for 30 minutes at 4 °C. The suspension was solubilised in 1% (w/v) n-dodecyl-β-D-maltopyranoside (DDM) supplemented with 0.1% cholesterol hemisuccinate (CHS) with 25 μM brain polar lipids extract (Avanti) for 45 minutes at 4 °C. Cell debris was removed by centrifugation at 140,000 × g, at 4 °C for 30 minutes.

TALON affinity resin (Cytiva) was added to the cleared lysate (1 mL resin per 10 g of cell pellet) in batch and incubated overnight at 4 °C, with stirring. The resin was washed with 10 column volumes (CV) of buffer containing 50 mM Tris-HCl pH 7.5, 150 mM NaCl, 0.05% (w/v) DDM, 0.005% (w/v) CHS, 2 mM adenosine 5′-triphosphate disodium salt hydrate (ATP), 10 mM MgCl_2_, and 30 mM imidazole pH 7.5. The resin was subsequently washed with 10 CV of buffer containing 50 mM Tris-HCl pH 7.5, 150 NaCl, and 30 mM imidazole supplemented with 0.1% (w/v) GDN, and with 10 CV of the same buffer supplemented with 0.05% (w/v) GDN. The protein was eluted in 50 mM Tris-HCl pH 7.5, 150 mM NaCl, 0.02% (w/v) GDN, and 200 mM imidazole pH 7.5. The protein was then treated with HRV-3C protease (in-house) for 3 hours at 4 °C to cleave the eGFP–twin-Strep-Tag-His tag, while being dialysed in buffer containing 50 mM Tris-HCl pH 7.5, 150 mM NaCl, and 0.006% (w/v) GDN.

The The eGFP–twin-Strep-Tag-His tag was removed by passing the sample through a nickel-nitrilotriacetic acid (Ni-NTA) HisTrap™ HP column (Cytiva) and impurities were removed by anion exchange chromatography with a pre-packed 1 mL HiTrap™ Q HP column (Cytiva). The protein was concentrated to 9 mg per mL, using an Amicon Ultra-15 centrifugal filter unit (Merck) with a 50,000 dalton cut-off, flash frozen in liquid nitrogen and stored at −80 °C.

On the day of grid preparation, the sample was further purified and buffer exchanged by size exclusion chromatography in buffer containing 50 mM Tris-HCl pH 7.5, 150 mM NaCl supplemented with either 10 mM GABA or 200 µM *SR*-THAP, using a Shodex kW803 column connected to an Äkta Go purifier system. Fractions were concentrated to 4 mg per mL and ultracentrifuged at 100,000 g for 30 minutes.

### Cryo-EM grid preparation and data collection

Prior to grid preparation, the protein sample was supplemented with 0.01% (w/v) of fluorinated octyl maltoside (Anatrace). HexAuFoil grids (Quantifoil) were glow discharged (Leica EM ACE200) at 15 mA for 180 s. 3 µl of sample was applied to the grids, blotted for 5 s with a blot force of +10 before being plunge-frozen in liquid ethane using a Vitrobot mark IV (ThermoFischer Scientific), at 100% humidity and 4°C in an environmental chamber.

Movies were collected in counting mode at 165,000× and 215,000× magnification (300 kV), yielding a raw pixel size of 0.729 Å and 0.571 Å per pixel, for *SR*-THAP-bound and GABA-bound sates, respectively, using EPU 3.6 software (ThermoFisher) on a Titan Krios G2 (ThermoFisher) equipped with a Selectris X energy filter set to a slit width of 10 eV and a Falcon 4i direct electron detector (ThermoFischer). A defocus range of –0.6 to –1.8 µm and a dose rate of around 10.2 electrons (e) per pixel per second were used, giving a total dose of 60 e^−^ Å^−2^ per movie.

### Cryo-EM image processing

The cryo-EM data sets were processed using cryoSPARC v4.5.3^60^. For *SR*-THAP-bound GAT3 a total of 44,460 cryo-EM movies in EER^61^ format were imported into cryoSPARC, motion corrected using patch motion correction with default parameters. The contrast transfer function (CTF) parameters were estimated using Patch CTF Estimation. Exposures with a CTF fit resolution lower than 3.6 Å were excluded during curation, resulting in 40,628 micrographs for further processing. Particles were picked by reference-free blob picker using a minimum and maximum particle diameter of 90 Å and 160 Å, respectively. A total of 9,151,454 particles were picked initially. Particles were extracted with a box size of 256 pixels and subjected to multiple rounds of 2D classification and ab initio-based 3D reconstruction to generate templates showing promising transmembrane helix features. 5,674,149 particles were then picked with the newly generated templates and extracted with a box size of 320 pixels (Supplementary Fig. 3). A single round of 2D classification was used to remove obvious bad classes, followed by multiples rounds of ab-initio in combination with heterogeneous refinement, yielding a map at 3.6 Å resolution with clearly defined 12 transmembrane helices and a ligand density from 492,706 particles. Using 3D classification, we could identify two sets of particle stack, one with a clear inhibitor density and one without. The corresponding two maps were further refined using 3D variability, 3D classification, non-uniform refinement^62^, and local refinement,, resulting in a 3D reconstruction of *SR*-THAP-bound GAT3 map at 2.9 Å resolution and substrate-free map at 3.9 Å resolution, based on a Fourier shell correlation (FSC) cutoff of 0.143 with 139,776 and 114,395 particles, respectively (Supplementary Fig. 3, Supplementary data table 1).

For the GABA-bound GAT3 a total of 33,686 cryo-EM movies in EER format were imported into cryoSPARC, and processed similar to *SR*-THAP-bound data set. Template were generated from the *SR*-THAP-bound GAT3, and GABA-bound GAT1 (EMD-33672). Combined with particles picked using reference-free blob picker, a total of 1,866,472 particles were extracted with a box size of 360 Å. Multiple rounds of 2D classification, ab initio, and heterogeneous refinement yielded a particle stack of 323,996 particles, of which a resulting map containing 199,550 particles showed clear density for 12 transmembrane helices and a density corresponding to the substrate GABA. Further local refinement, 3D classification and non-uniform refinement resulted in a 3D reconstruction map at 3.18 Å with 95,516 particles (Supplementary Fig. 4, Supplementary data table 1).

### Model building and refinement

The initial GAT3 model was was derived using AlphaFold2^63^ (AF-P48066-F1-v4). The predicted model was manually fitted into the cryo-EM map in UCSF ChimeraX^64^ v.1.8 and was initially refined using Namdinator^65^. The models were refined using Phenix^66^ v1.21.1-5286 followed by visual examination and manual rebuilding in Coot^67^ v0.9.8.6 and ISOLDE^68^ v1.6. The ligand-restraint files for refinement were generated by phenix.elbow. Figures were prepared using ChimeraX^68^ v1.8.

The final model of *SR*-THAP-bound lacks the initial 54 residues, residues 188–192 in EL2, 248-251 in TM5, and the last 32 residues of the C terminus due to high flexibility and poor density of the regions. The final model of GABA-bound lacks the initial 50 residues, residues 187–194 in EL2, 246-253 in TM5, residues 431-434, and the last 33 residues of the C terminus due to high flexibility and poor density of the regions. The final model of substrate-free GAT3 lacks the initial 56 residues, residues 187–195 in EL2, 247-251 in TM5, and the last 32 residues of the C terminus. Refinement and validation statistics are presented in Supplementary data Table 1.

### HEK293T cell culture and transfection

HEK293T cells were cultured in DMEM containing GlutaMAX-I (Gibco, Thermo Fisher Scientific), supplemented with 10% fetal bovine serum (FBS) (Gibco) and 1% penicillin/streptomycin (P/S) (Invitrogen, Thermo Fischer Scientific), and maintained at 37 °C in a humidified incubator with 5% CO_2_. At approximately 60% confluency, cells were transfected in 10 cm dishes with 4 μg of plasmid, using PolyFect reagent according to the manufacturer’s protocol (Qiagen), but with half the recommended PolyFect volume. Approximately 20 hours post-transfection, cells were plated in white 96-well polystyrene microplates (CulturPlate, PerkinElmer) coated with poly-D-lysine (PDL; Sigma-Aldrich) at a density of 50,000 cells/well and left overnight (37 °C and 5% CO2). The radioligand competition uptake assay was performed approximately 48 hours post-transfection.

### GABA uptake experiments assay

The [^3^H]GABA and [^3^H]glycine competition uptake assays were performed as previously described^17^ using [2,3-3H(N)]GABA (specific activity 35 Ci/mmol) and 3H-glycine (specific activity of 42.6 Ci/mmol), both from PerkinElmer. Briefly, cells were washed with 100 µL room temperature assay buffer (HBSS supplemented with 20 mM HEPES, 1 mM CaCl_2_,1 mM MgCl_2_, pH 7.4) before incubation with 75 µL assay buffer containing 30 nM radioligand ([^3^H]GABA or [^3^H]Glycine), with or without concentration series of test compounds for 3 min at 37 °C. This was followed by washing of plate with 3 × 100 µL ice-cold assay buffer, before addition of 150 µl Microscint™ 20 scintillation liquid (PerkinElmer) to each well. The plate was then shaken for at least one hour and counted in a TopCount NXT Microplate Scintillation & Luminescence Counter (PerkinElmer). Full uptake was determined with plain radioligand solution, while full inhibition was assessed in the presence of 3 mM GABA or 3 mM glycine in the radioligand solution. For preincubation experiments, cells were incubated with in plain assay buffer or a concentration series of test compounds in assay buffer for 2 hours at 37 °C. Following preincubation, wells were washed with 100 µL of room-temperature assay buffer before proceeding with the radioligand competition uptake assay as described above. All concentration response curves were performed in triplicate in at least three independent experiments.

### Pharmacological data analysis

Data were collected from at least three biologically independent experiments (from transfection of cells) and individual data points represent the mean of triplicate measurements. All data were normalized to full inhibition assessed in the presence of either 3 mM GABA or 3 mM glycine. Data was analyzed using GraphPad Prism 10.3 (GraphPad Software, San Diego, CA, USA). The IC_50_ values were determined by fitting to a nonlinear regression model of log([inhibitor]) versus normalized response with variable slope: *y* = 100 / (10^Hillslope•log(IC50-x)^ + 1)

### Molecular docking

Compounds *RS*-THAP, DPPM-1457 and (*S*)-SNAP-5114 were prepared in Maestro using the 2D sketch editor and were prepared for docking with Ligprep with default settings (Schrödinger Release 2024-2: Maestro, Ligprep, Schrödinger, LLC, New York, NY, 2024). As at pH 7 the compounds are zwitterions, they are deprotonated at the carboxylic acid and protonated at the basic tertiary nitrogen. Consequently, for each compound, two stereoisomers with a different absolute configuration at the ammonium nitrogen were generated, but only compounds with an absolute configuration at the nitrogen comparable to that of tiagabine in GAT1 (PDB IDs 7Y7Z and 7SK2) ^23,35^ and of *SR*-THAP in GAT3 (current study) were retained. GAT3 was imported in maestro as a pdb file, preprocessed for docking and H-bonds optimized using protein preparation workflow^69^. Flexible docking without post-docking minimization was performed using extra precision (XP) Glide with default settings without post-docking minimization, and the best scored pose according to the emodel score was selected as representative for each compound^70^.

