## Supplementary Figures and Supplementary data Table 1 for "Structural basis for selective inhibition of human GABA transporter GAT3"

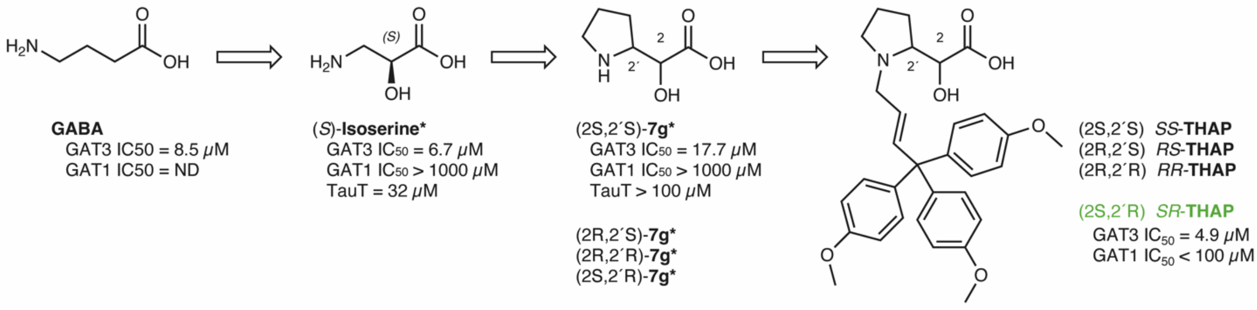


**Supplementary Fig. 1 Design strategy leading to *SR*-THAP and its stereoisomers.** *Data for (*S*)-isoserine and for analogues 7g, as well as naming of 7g are taken from ref. 17. Where an $IC_{50}$ for GAT3 is not specified, the compound is inactive ($IC_{50}$ > 100 μM).

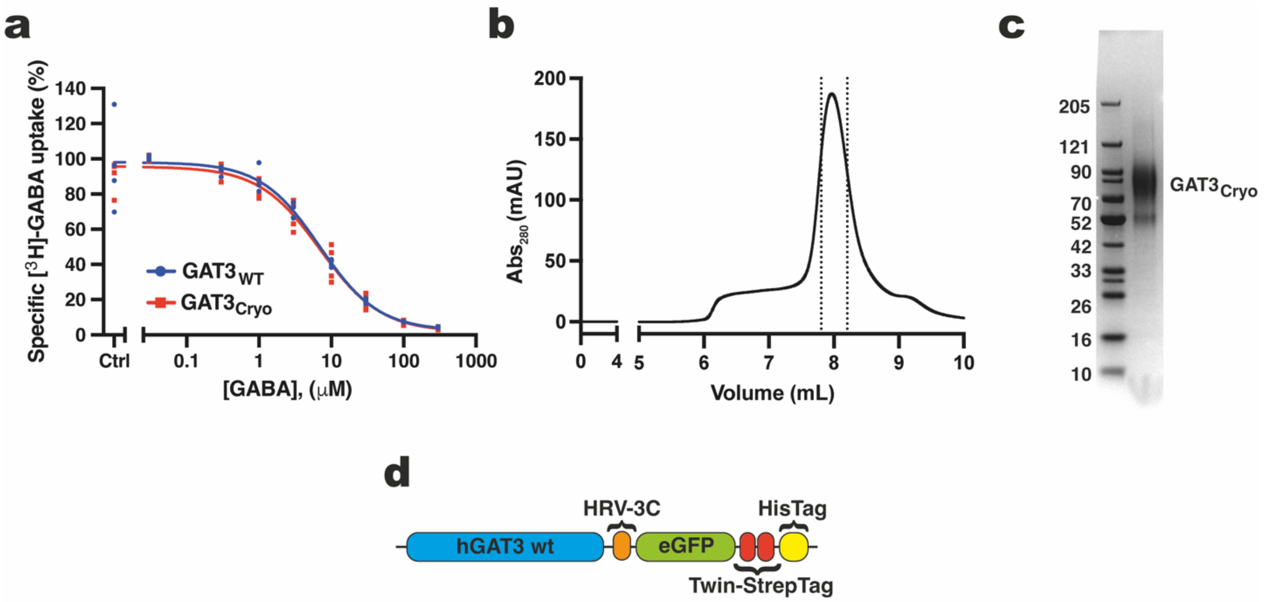


**Supplementary Fig. 2 | Characterization and purification of GAT3$_{Cryo}$ construct. a,** Measurement of GABA inhibition of [$^3$H]GABA uptake into HEK293T cells expressing GAT3$_{WT}$ and GAT3$_{Cryo}$ with IC$_{50}$ values of 6.9 [4.9; 10] μM and 7.0 [5.3; 9.3] μM (no difference, $p = 0.968$). Experiment was performed as four independent replicates, each measured as triplicate measurements ($n = 4$). **b,** Size exclusion chromatography of GAT3$_{Cryo}$ on a Shodex protein KW-803 HPLC column, tracking the absorbance at 280 nm, with indicated vertical dotted lines of the fractions used for grid preparation. **c,** Coomassie stained SDS-PAGE gel showing hGAT3 after HRV-3C cleavege of affinity taggs. **d,** Illustration of the elements in the GAT3$_{Cryo}$ construct, including a HRV-3C cleavege site, enhanced GFP (eGFP), a Twin-Strep affinity tag, and a decahistidine tag.

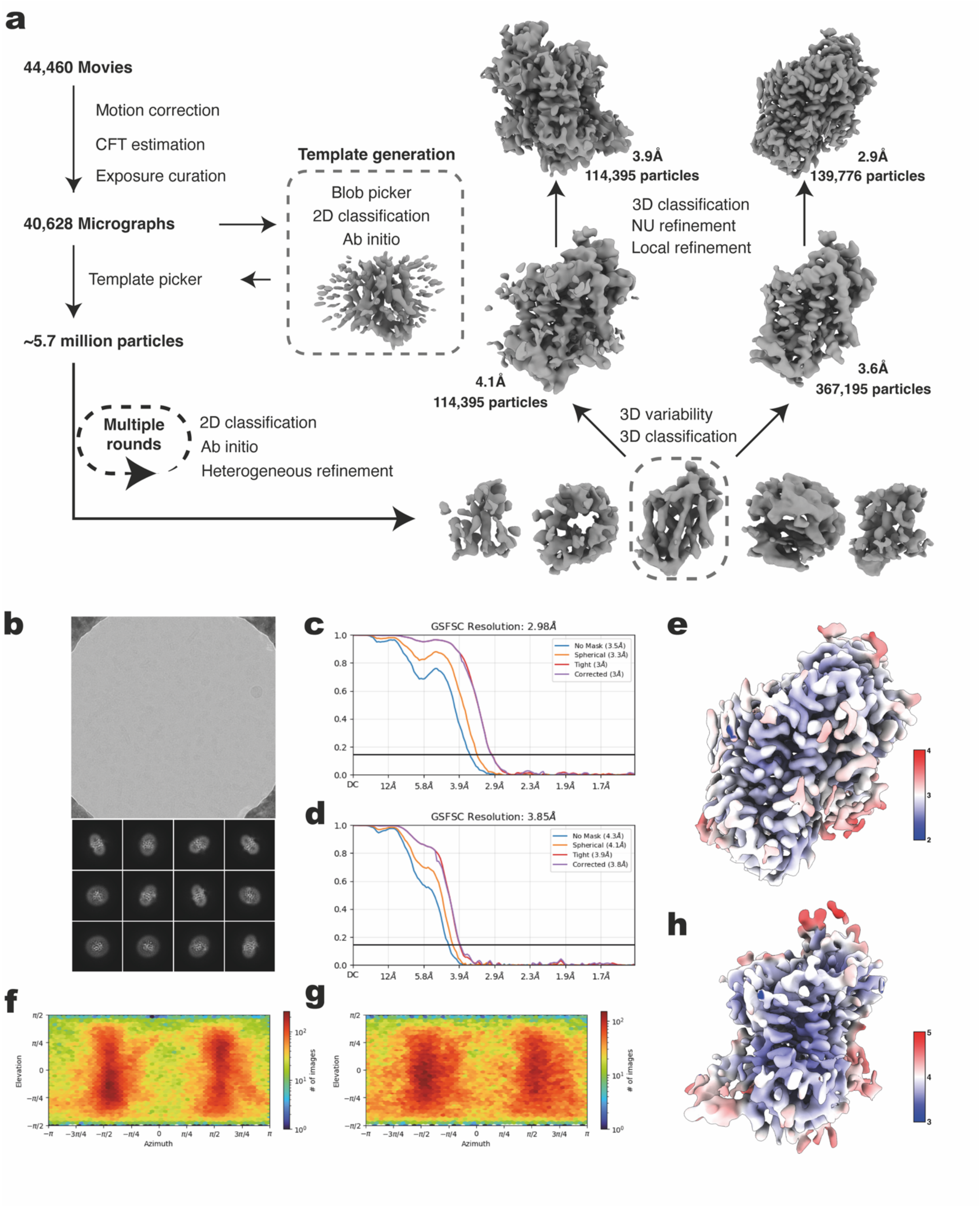


**Supplementary Fig. 3 | Cryo-EM data processing workflow of *SR*-THAP-bound GAT3. a,** Cryo-EM workflow. In brief, 44,460 movies were selected after manual curation. 7,5377,655 particles were initially picked from which 106,582 particles were selected to generate 2 ab initio models. The best model was then used as a template to pick particles. Following a 2D classification, selected particles were used to generate 5 ab initio classes and after two rounds of heterogenous refinement and a 3D classification, a 3.6 Å map was generated with a clear density for *SR*-THAP and a 4.1 Å map without that density was generated. The final 2.9 Å map for the *SR*-THAP bound, and 3.9 Å map for the substrate-free were generated after several rounds of non-uniform refinement, 2D classification, a reference-based motion correction, and a final non-uniform refinement. **b**, Representative motion-corrected micrograph and representative 2D class averages. **c,** The gold standard Fourier shell correlation (GSFSC) curve with FSC= 0.143 cut-off represented by a black horizontal line. **d**, The conical FSC curve. **e**, Angular sampling of the final reconstruction. **f**, Local resolution map.

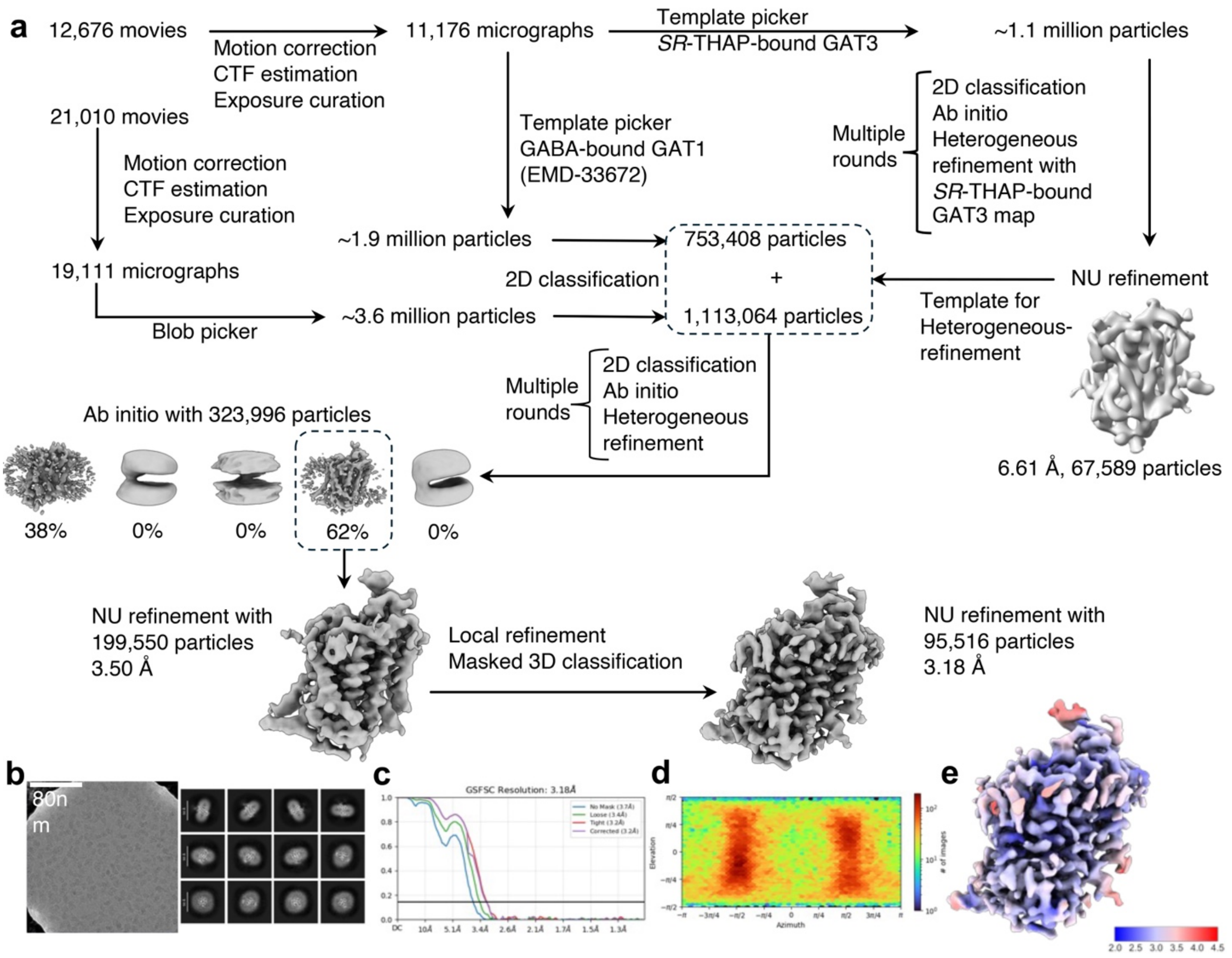


**Supplementary Fig. 4 | Cryo-EM data processing and analysis of GABA-bound GAT3.**

**a,** Cryo-EM workflow. In brief, 11,176 movies were selected after manual curation of the first data set. ~1.1 million particles were initially picked with a template using a CA79M2-bound GAT3 map. After several rounds of 2D classification, Ab initio reconstruction and Heterogeneous refinement 67,589 particles were refined with a NU refinement to 6.61 Å resolution. After a template pick with a GABA-bound GAT1 map (EMD-33672), ~1.9 million picked particles were classified in 2D, leaving 753,408 particles after selection. After manual curation of the second dataset, ~3.6 million particles were picked with a blob picker from 19,111 micrographs and 2D classified down to 1,113,064 particles. The selected particles originating from the GABA-bound GAT1 template pick (data set 1) and blob picker (data set 2) were combined in a Heterogeneous refinement using the first NU refinement as an initial template. After multiple rounds of 2D classification, Ab initio reconstruction and Heterogeneous refinement 199,550 particles were selected from a 5 class Ab initio reconstruction and refined to 3.50 Å with a NU refinement. Subsequent Local refinement and masked 3D classification resulted in the final NU refinement of 95,516 particles at 3.18 Å resolution. **b**, Representative motion-corrected micrograph image and representative 2D classification averages. **c**, The gold standard Fourier shell correlation curve (GSFSC) of the final NU refinement. The FSC = 0.143 cut-off is represented by a black horizontal line. **d,** Angular sampling of the final reconstruction. **e,** Local resolution map with a FSC = 0.143 cut-off.

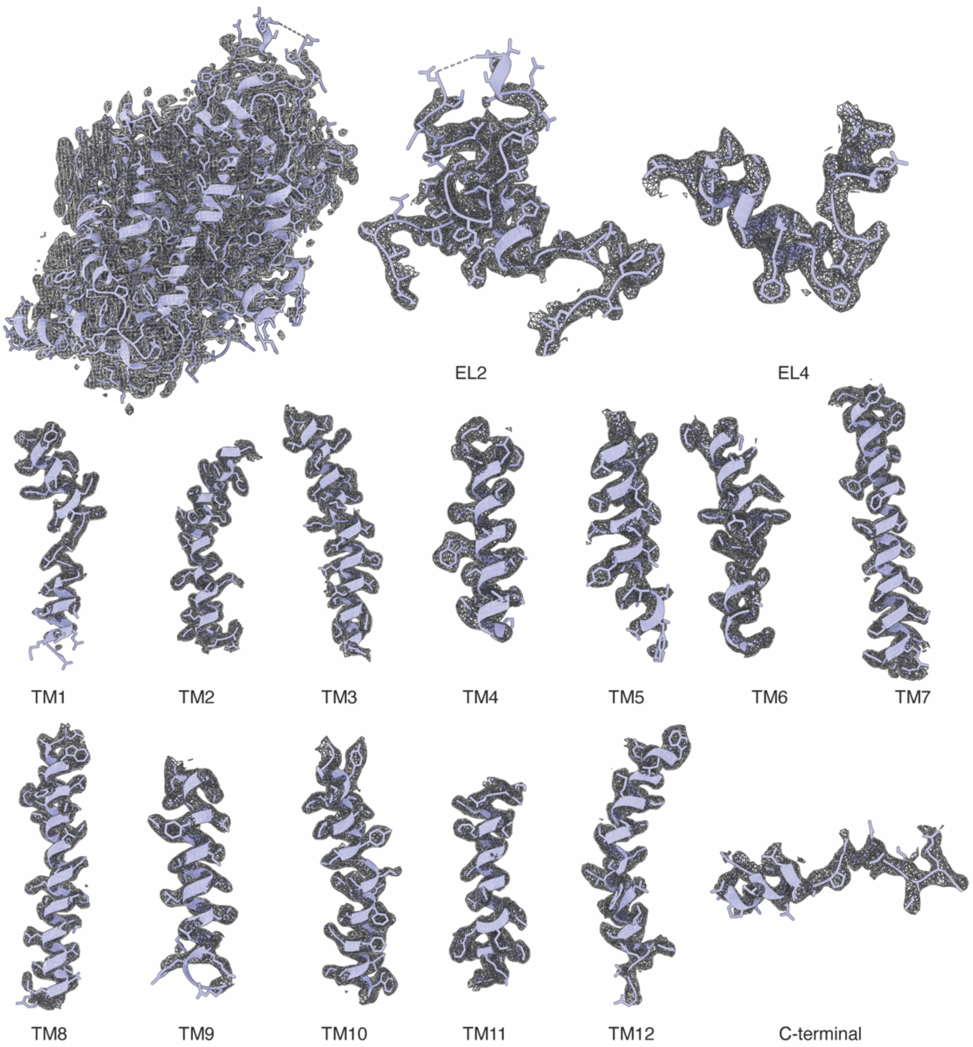


**Supplementary Fig. 5 | Cryo-EM density of *SR*-THAP-bound GAT3**. The overall structure of *SR*-THAP-bound GAT3 (top left, PDB ID 9QO8) and magnified density views of transmembrane helices (contour level = 0.072 in ChimeraX), extracellular loop (EL) 2 (with the conserved C171 – C180 disulfide bridge), as well as EL4 and the C terminus are shown.

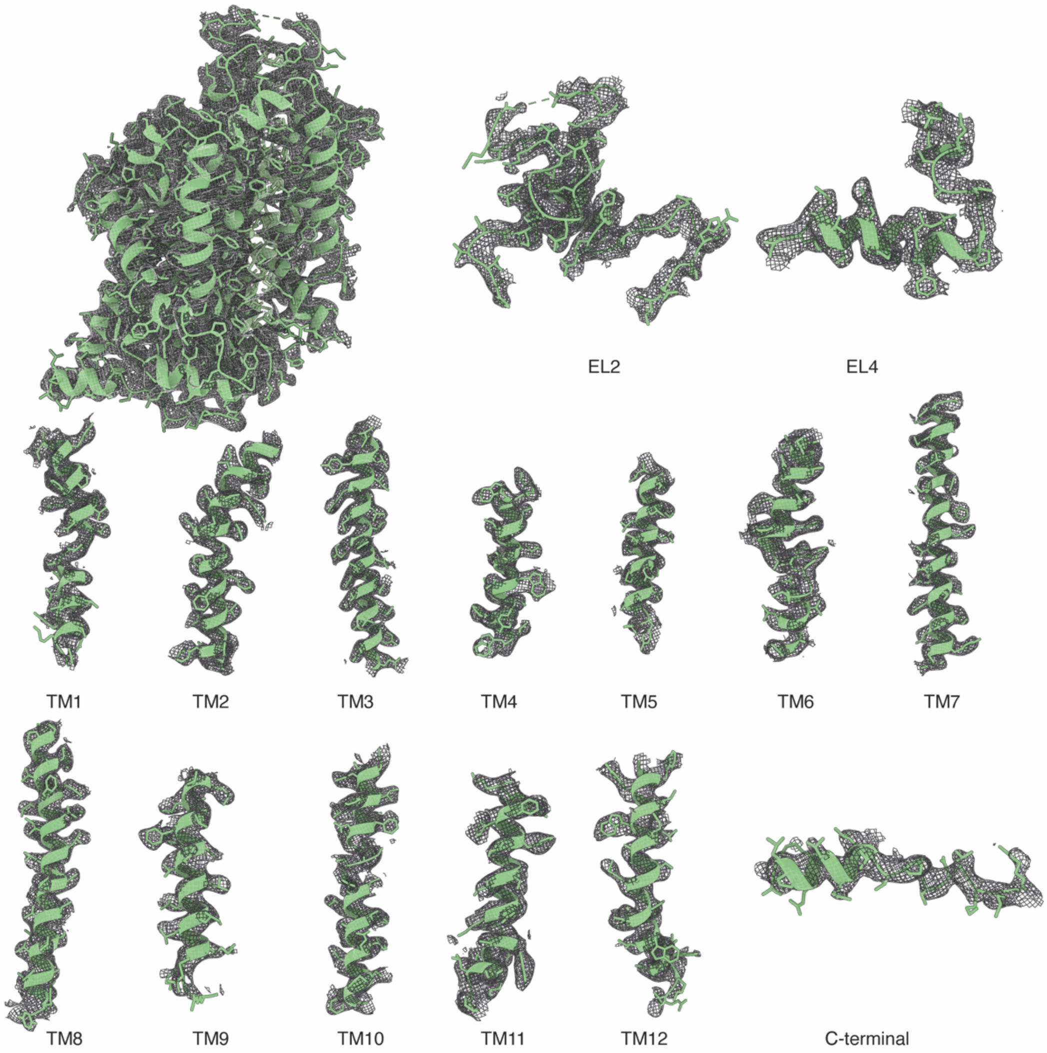


**Supplementary Fig. 6 | Cryo-EM density of GABA-bound GAT3**. The overall structure of GABA-bound GAT3 (top left, PDB ID 9QO9) and magnified density views of transmembrane helices (contour level = 0.055 in ChimeraX corresponding to 5 σ), EL2 (with the conserved C171 and C180 residues), as well as EL4 and the C terminus are shown.

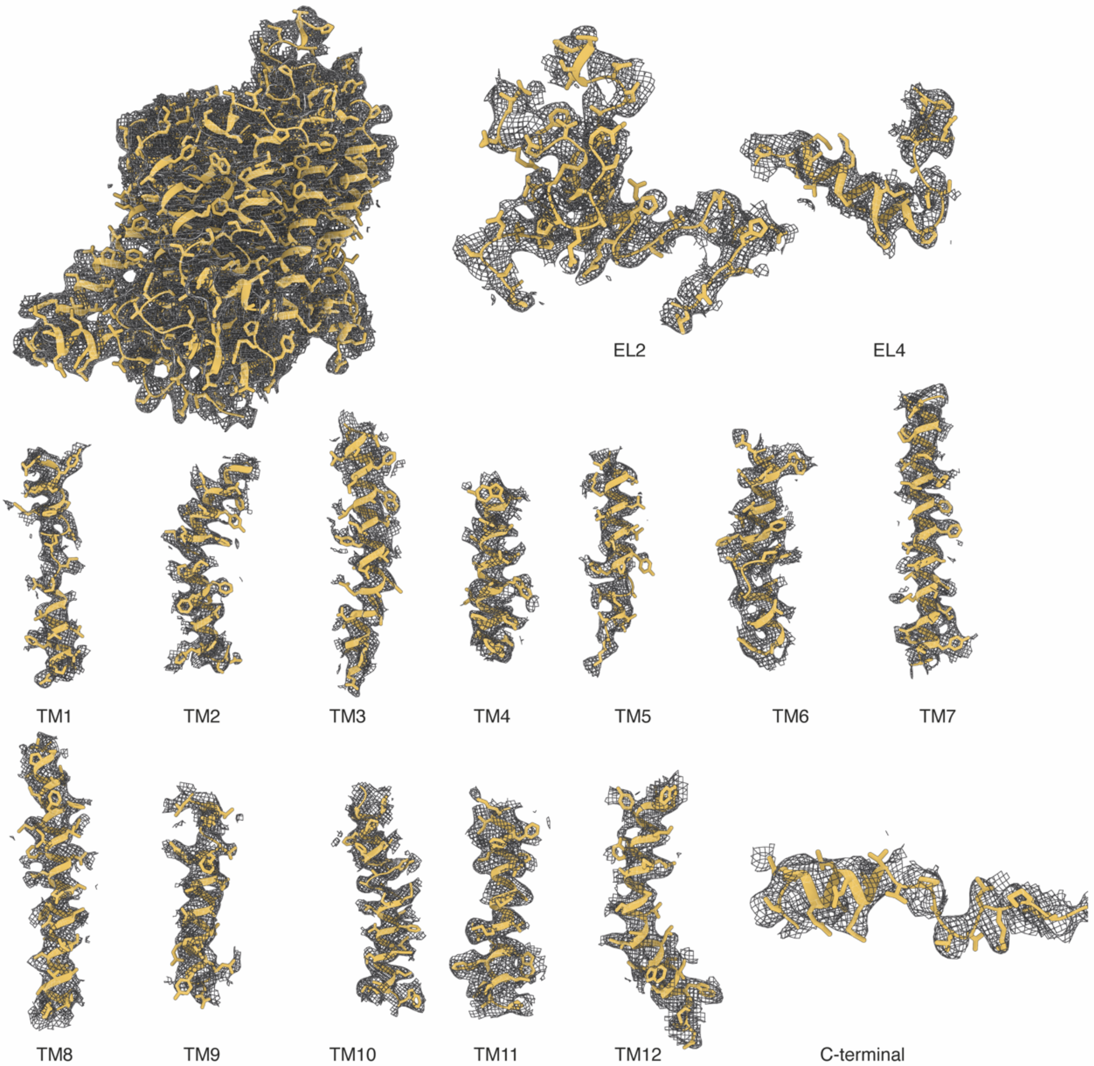


**Supplementary Fig. 7 | Cryo-EM density of apo state GAT3**. The overall structure of the unbound GAT3 (top left, PDB ID 9QOA) and magnified density views of transmembrane helices (contour level = 0.036 in ChimeraX corresponding to 5 r.m.s.d), extracellular loop (EL) 2 (with the conserved C171 – C180 disulfide bridge), as well as EL4 and the C terminus are shown.

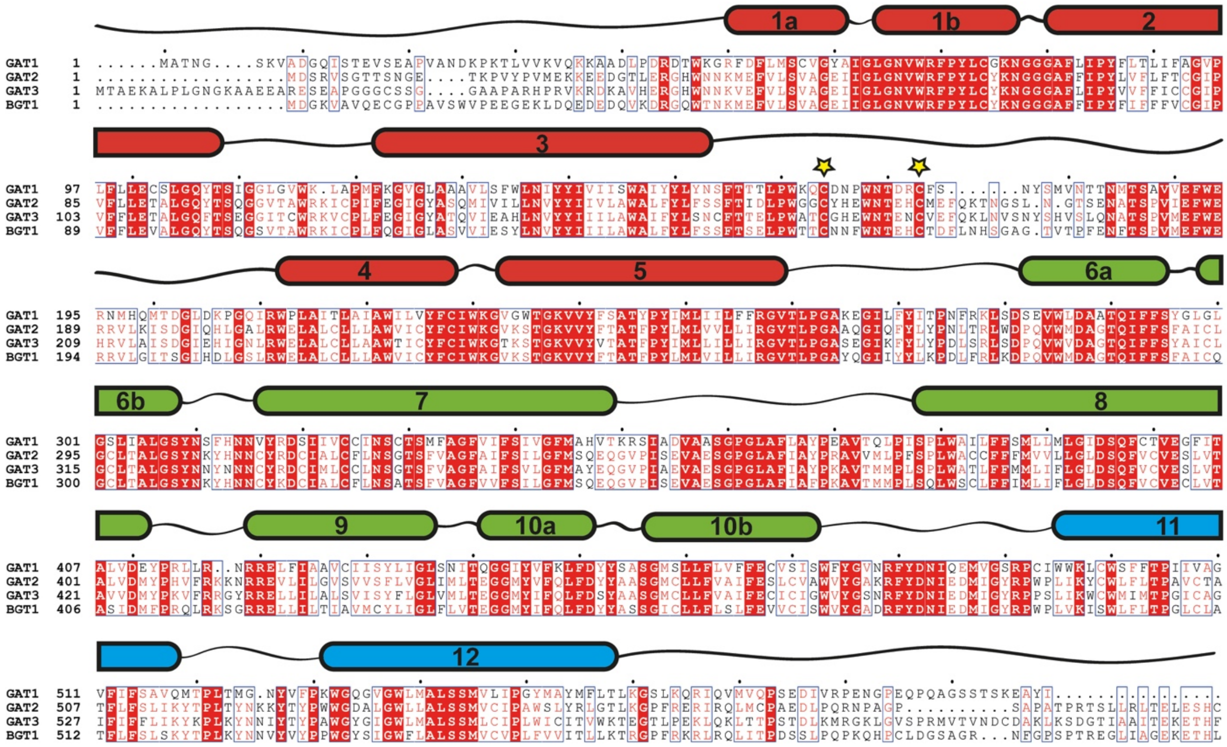

**Supplementary Fig. 8 | Sequence alignment of the four human GABA transporters.** The transmembrane segments are annotated and coloured according to the pseudo-symmetric fold (TM1-5 – red; TM6-10 – green; TM11-12 – blue). Yellow stars indicate conserved cysteine residues in EL2 (C171 – C180 in GAT3).

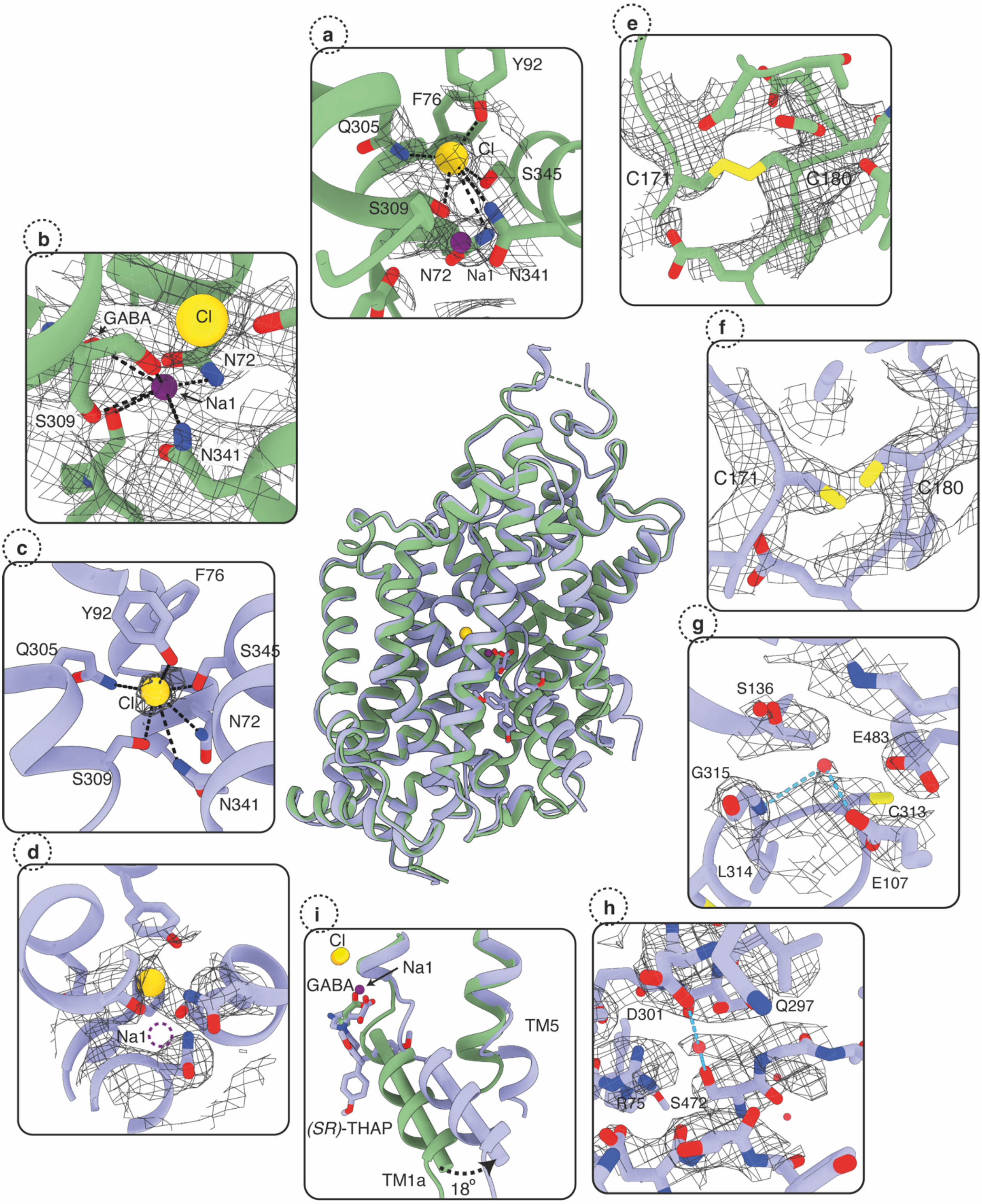


**Supplementary Fig. 9 | Ion coordination, structural water molecules, the conserved disulfide bridge, and angular movement of TM1a.** Structural overview of *SR*-THAP-, and GABA-bound hGAT3 (middle). **a, b,** Chloride and sodium binding sites identified by clear densities in the GABA-bound GAT3, with indicated interactions as black dashed lines. **c, d,** Chloride site and empty sodium site and densities in the *SR*-THAP-bound GAT3, with indicated interactions shown as black dashed lines. **e, f,** Conserved disulfide bridge between C171 and C180, was not observed in the GABA-bound GAT3, but was clearly defined in the *SR*-THAP-bound GAT3. **g, h,** Structural water molecules with clearly defined densities in *SR*-THAP-bound GAT3, were observed in the coordinating of the conserved CLG motif of the unwound part of TM6 (**g**), as well as coordinating the extracellular gating residues R75, S472, and D301 (**h**). **i,** An outward shift of

TM1a between GABA-, and *SR*-THAP-bound GAT3 of 18º is consistent with the opening of the intracellular permeation pathway. The *SR*-THAP-bound GAT3 structure is shown as cartoon in light blue with corresponding densities at contour level of 0.072 (in ChimeraX), and GABA-bound GAT3 structure is shown as cartoon in green with corresponding densities at contour level of 0.055 (in ChimeraX). Chloride ions are shown as yellow spheres, sodium purple and water molecules as red spheres.

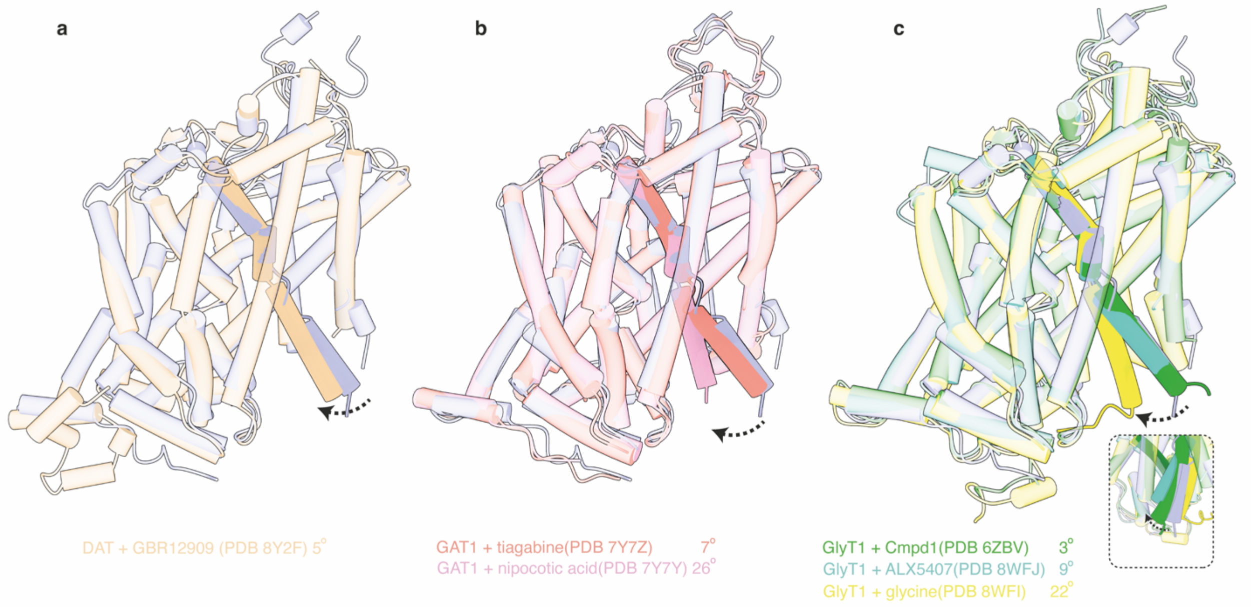


**Supplementary Fig. 10 | Measuring TM1 angular movement between *SR*-THAP-bound hGAT3 structure and other SLC6 members**. Shift of TM1 was measured between the Cα-Cα atoms of the first residue on the N terminus of the TM1a as the moving points and G69 in GAT3 as the fixed point. The overlay of *SR*-THAP-bound GAT3 (in blue) with GBR12909-bound hDAT (light brown) (**a**), with tiagabine-bound (pink) and nipecotic acid-bound (red) hGAT1 (**b**) and Cmpd1-bound (green), ALX5407-bound (cyan), and glycine-bound (yellow) hGlyT1 (**c**) are shown.

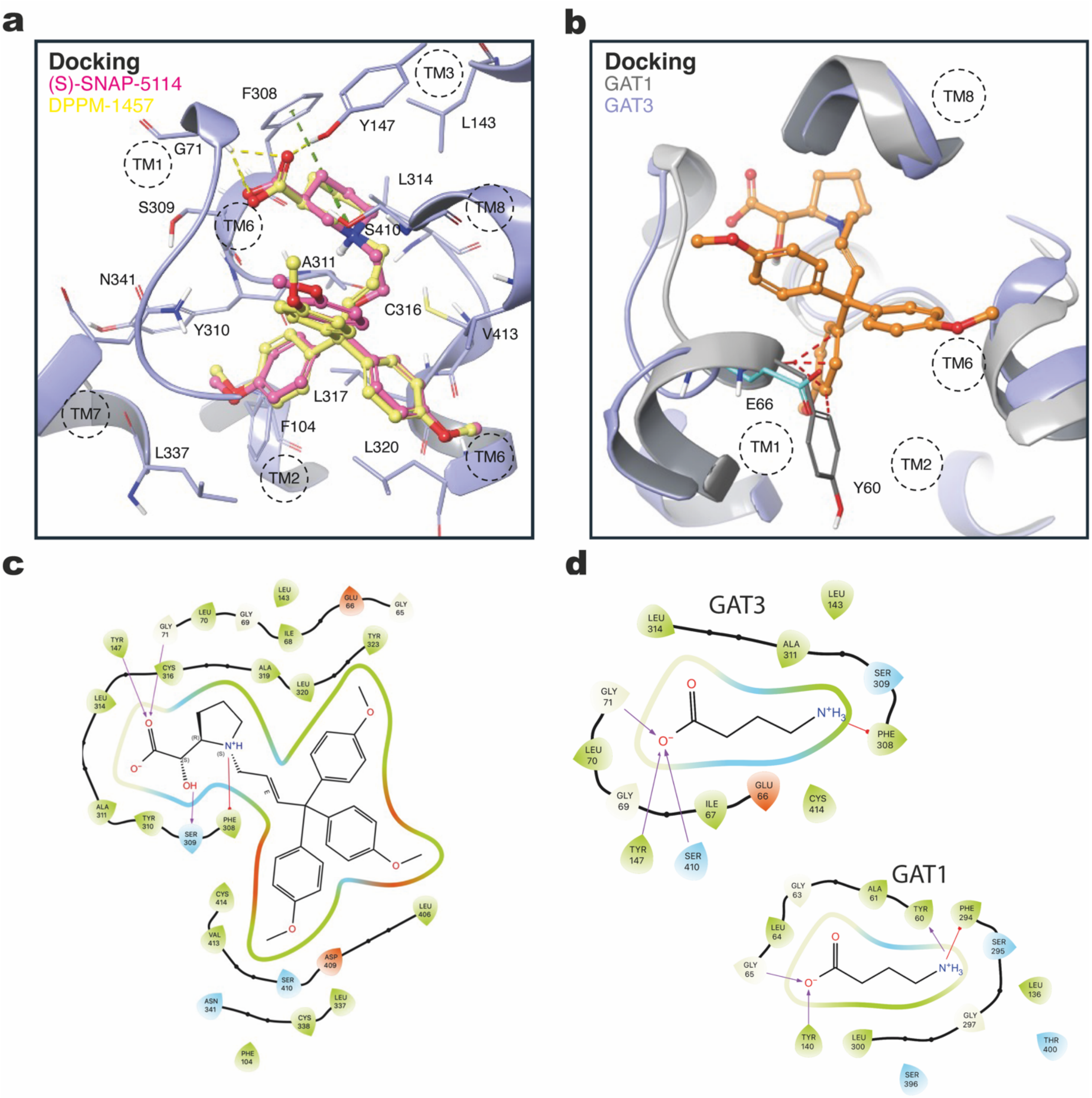


**Supplementary Fig. 11 | a,** Close-up view of the docking pose of (S)-SNAP-5114 (pink) and of DPPM-1457 (yellow) in GAT3 (ribbons as azure cartoons). Interacting residues within 3 Å are displayed in grey. H-bonds and cation-$\pi$ interactions are displayed with yellow and green dashed lines, respectively. **b,** Overlay between *SR*-THAP-bound GAT3 (ribbons as azure cartoons) and tiagabine-bound GAT1 (PDB ID: 7Y7Z, ribbons as grey cartoons) illustrating the steric clashes between Y60 and *SR*-THAP. **c,** Schematic illustration of the binding mode of *SR*-THAP at GAT3. Residues within 3 Å are displayed. Hydrogen bonds are shown as pink arrows, and $\pi$-cation interactions as red lines.

**Supplementary Data Table 1 | Cryo-EM data collection, refinement and validation statistics.**

| | *SR*-THAP-bound GAT3 (EMD-53259) (PDB ID 9QO8) | GABA-bound GAT3 (EMDB-53260) (PDB ID 9QO9) | Substrate-free GAT3 (EMDB-53261) (PDB ID 9QOA) |
|---|---|---|---|
| **Data collection and processing** | | | |
| Magnification | 165,000× | 215,000× | 165,000× |
| Voltage (kV) | 300 | 300 | 300 |
| Electron exposure ($e^{-}$/Å$^2$) | 60 | 60 | 60 |
| Defocus range (μm) | –0.6 to –1.8 | –0.6 to –1.8 | –0.6 to –1.8 |
| Pixel size (Å) | 0.729 | 0.571 | 0.729 |
| Symmetry imposed | C1 | C1 | C1 |
| Initial particle images (no.) | 9,179,048 | 5,482,177 | 9,179,048 |
| Final particle images (no.) | 139,776 | 96,516 | 114,395 |
| Map resolution (Å) | 2.90 | 3.18 | 3.97 |
| FSC threshold | 0.143 | 0.143 | 0.143 |
| Map resolution range (Å) | 2.0 – 4.0 | 2.0 – 4.5 | 3.0 – 5.0 |
| **Refinement** | | | |
| Initial model used (PDB code) | AF-P48066-F1 | AF-P48066-F1 | 9QO8 |
| Model resolution (Å) | 2.70 | 3.10 | 3.99 |
| FSC threshold | 0.143 | 0.143 | 0.143 |
| Map sharpening *B* factor (Å$^2$) | | | |
| Model composition | | | |
| Non-hydrogen atoms | 4298 | 4204 | 4256 |
| Protein residues | 535 | 527 | 535 |
| Ligands | 1 | 1 | 0 |
| Ions | 1 | 2 | 0 |
| Waters | 6 | 2 | 0 |
| *B* factors (Å$^2$) | | | |
| Protein | 60.47 | 84.06 | 143.82 |
| Ligand | 62.66 | 56.48 | NA |
| R.m.s. deviations | | | |
| Bond lengths (Å) | 0.003 (0) | 0.004 (0) | 0.005 (0) |
| Bond angles (°) | 0.698 (1) | 0.843 (0) | 1.053 (4) |
| Validation | | | |
| MolProbity score | 1.15 | 1.49 | 2.31 |
| Clashscore | 3.63 | 7.05 | 13.66 |
| Poor rotamers (%) | 0.44 | 0.89 | 0.42 |
| CaBLAM outliers (%) | 0.57 | 0.98 | 1.72 |
| Ramachandran plot | | | |
| Favored (%) | 98.87 | 97.50 | 93.19 |
| Allowed (%) | 1.13 | 2.50 | 6.62 |
| Disallowed (%) | 0.00 | 0.00 | 0.19 |
